# A targetable PREX2/RAC1/PI3Kβ signalling axis confers resistance to clinically relevant therapeutic approaches in melanoma

**DOI:** 10.1101/2024.02.26.580534

**Authors:** Catriona A. Ford, Dana Koludrovic, Patricia P. Centeno, Mona Foth, Elpida Tsonou, Nikola Vlahov, Nathalie Sphyris, Kathryn Gilroy, Courtney Bull, Colin Nixon, Bryan Serrels, Alison F. Munro, John C. Dawson, Neil O. Carragher, Valeria Pavet, David C. Hornigold, Philip D. Dunne, Julian Downward, Heidi C. E. Welch, Simon T. Barry, Owen J. Sansom, Andrew D. Campbell

## Abstract

Metastatic melanoma remains a major clinical challenge. Large-scale genomic sequencing of melanoma has identified *bona fide* activating mutations in *RAC1*, with mutations of its upstream regulator, the RAC-GEF *PREX2*, also commonly detected. Crucially, *RAC1* mutations are associated with resistance to BRAF-targeting therapies. Despite the role of its homologue PREX1 in melanomagenesis, and evidence that some truncating *PREX2* mutations drive increased RAC1 activity, no hotspot mutations have been identified, and the impact of *PREX2* mutation remains contentious. Here, we use genetically engineered mouse models and patient-derived BRAFV600E-driven melanoma cell lines to dissect the role of PREX2 in melanomagenesis and response to therapy. We show that while PREX2 is dispensable for the initiation and progression of melanoma, its loss confers sensitivity to clinically relevant therapeutics. Importantly, genetic and pharmacological targeting of the RAC1 effector kinase PI3Kβ phenocopies *PREX2* loss, sensitizing our model systems to therapy. Our data reveal a druggable PREX2/RAC1/PI3Kβ signalling axis in *BRAF*-mutant melanoma that could be exploited clinically.

**Statement of Significance:** Metastatic melanoma remains both a clinical problem, and an opportunity for therapeutic benefit. Co-targeting of the MAPK pathway and the PREX2/RAC1/PI3Kβ has remarkable efficacy and outperforms monotherapy MAPK targeting *in vivo*.

## Introduction

Despite the development of efficacious targeted therapeutic approaches, resulting in improved overall survival rates over the last 20 years, metastatic melanoma remains a clinical problem. While surgery is often curative in early-stage disease, the prognosis for patients diagnosed with metastatic melanoma remains poor, due to the ineffectiveness of surgery in disseminated disease alongside rapidly acquired resistance to targeted therapies. Improved understanding of the molecular basis of melanoma, and of both the response and resistance to targeted therapy, remains critical.

Key to success of targeted therapeutics for melanoma are two concurrent, yet independent, approaches – effective targeting of tumour cell-intrinsic driver mutations and their effector pathways, and reversal of tumour-driven immune suppression. Mutations in *BRAF* occur in ∼50% of melanomas, with the vast majority introducing a phosphomimetic V600E substitution (*BRAF*V600E) that confers constitutive kinase activity^1^. While these oncogenic mutations commonly occur in benign melanocytic precursor lesions (naevi), subsequent accumulation of oncogenic mutations or loss of tumour-suppressor genes, such as *CDKN2A*, *PTEN*, or *TP53,* ultimately drives progression to melanoma. Indeed, oncogenic mutations in *BRAF* frequently co-occur with inactivation of the tumour suppressor PTEN in melanomas^2,3^, eliciting aberrant activation of the pro-proliferative MAPK pathway and PI3K/AKT/mTOR signalling, respectively. PTEN deficiency cooperates with BRAFV600E, contributing to melanomagenesis^4^, resistance to BRAF inhibition^4^, and metastasis^5^. In addition, most melanomas typically harbour a mutational signature driven by ultraviolet-(UV)-radiation exposure^6^, although this is absent in ∼15% of cases^7^.

The prevalence of *BRAF* mutations, paucity of effective clinical approaches for late-stage disease, and poor prognosis made *BRAF*-mutant metastatic ideal for early adoption of novel small molecules such as vemurafenib, which selectively targets the BRAFV600E oncoprotein^8^. While targeted BRAF inhibition with vemurafenib or dabrafenib, or combined BRAF and MEK1/2 inhibitors (vemurafenib/trametinib, dabrafenib/cobimetinib or encorafenib/binimetinib) are now mainstays of the management of metastatic melanoma, the emergence of therapeutic resistance poses a significant clinical challenge^9^, and calls for new strategies targeting therapeutic resistance, melanoma recurrence, and metastatic progression (Fig 1A).

**Fig. 1:**
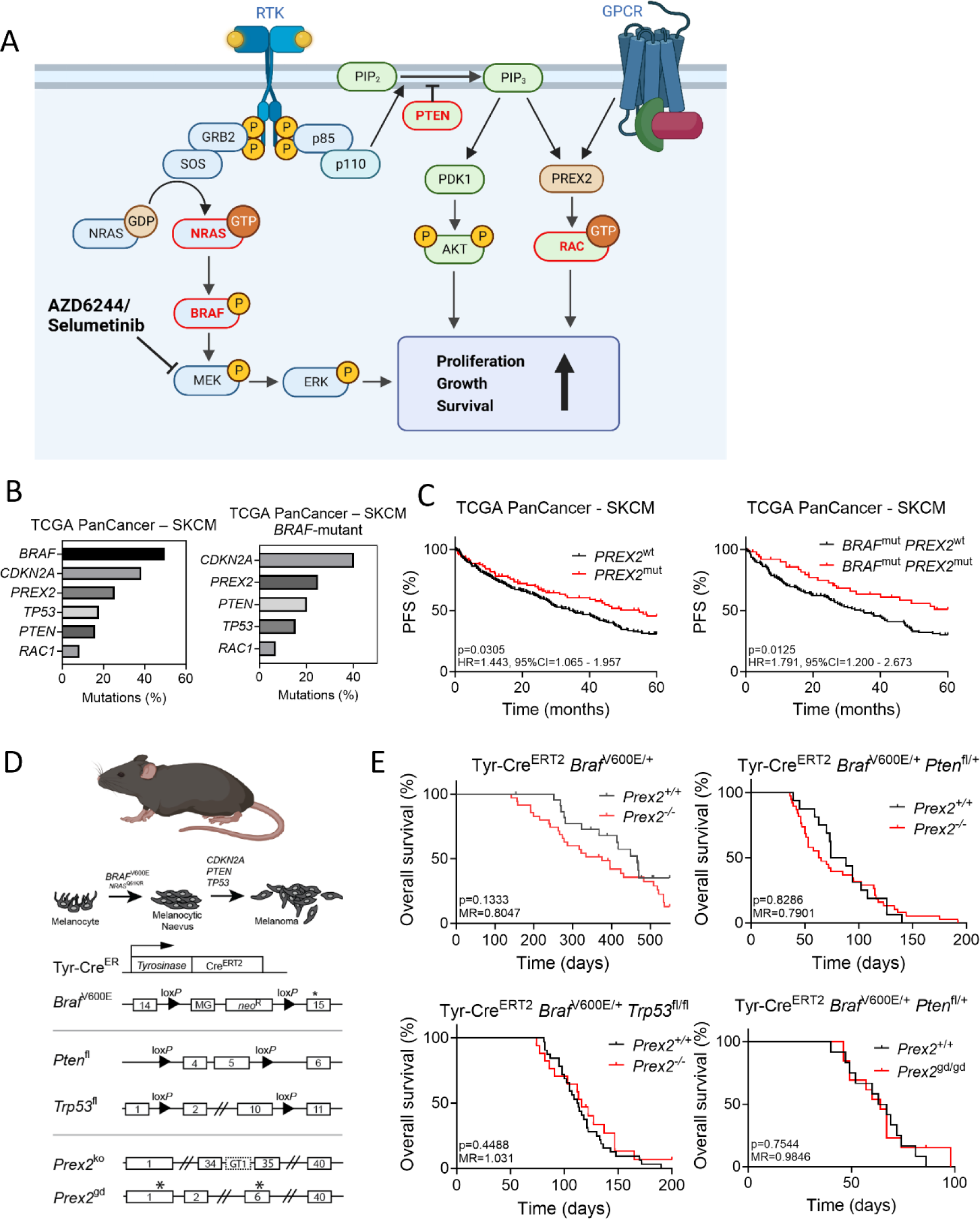
PREX2 ablation does not impact melanoma progression *in vivo*. A) Schematic of receptor tyrosine kinase (RTK) and G-protein–coupled receptor (GPCR) signalling in melanoma. Arrows represent activation, blunt-ended lines represent inhibition, and circled P represents phosphorylation. Schematic created with Biorender.com. B) Histograms of mutation frequencies of select commonly mutated genes in total (left) and *BRAF*-mutant (right) skin cutaneous melanoma (SKCM) cohorts from The Cancer Genome Atlas (TCGA) PanCancer study, accessed via cBioPortal on 23/06/2023. C) Progression-free survival (PFS) of *PREX2* mutant *vs* wild-type cases in the curated cohort of SKCM patients from the TCGA PanCancer cohort, censored at 5 years. Left panel, all cases (n=381; *PREX2* wild-type = 290, *PREX2* mutant = 91); Right panel, *BRAF*-mutant cases (n=207; *PREX2* wild-type = 155, *PREX2* mutant = 52). D) Top panel, schematic timeline of melanoma development (melanocytic dysplasia, naevus formation, and progression to melanoma) with common clinically relevant mutations indicated. Bottom panels, schematics of transgenic alleles used to generate melanoma GEMMs. Exons are boxed, with exon numbers indicated. Asterisks indicate the location of mutations. loxP sites, filled triangles; MG, minigene encoding exons 15–18 of wild-type *Braf* with a 5′ splice acceptor; neo^R^, neomycin phosphotransferase gene; GT1, gene trap vector containing the β-galactosidase/neomycin-resistance fusion gene (β-geo); *Prex2*^gd^, *Prex2*^E22A,N204A^ allele. E) Top left panel, overall survival of Tyr-Cre^ERT2^ *Braf*^V600E/+^ *Prex2*^+/+^ (n=23; median survival, 466 days) *vs* Tyr-Cre^ERT2^*Braf*^V600E/+^ *Prex2*^-/-^ (n=35; median survival, 375 days) mice, *p*=0.1333. Top right panel, overall survival of Tyr-Cre^ERT2^ *Braf*^V600E/+^ *Pten*^fl/+^ *Prex2*^+/+^ (n=16; median survival, 81 days) *vs* Tyr-Cre^ERT2^ *Braf*^V600E/+^ *Pten*^fl/+^ *Prex2*^-/-^ (n=38; median survival, 64 days) mice, *p*=0.8286. Bottom left panel, overall survival of Tyr-Cre^ERT2^ *Braf*^V600E/+^ *Trp53*^fl/fl^ *Prex2*^+/+^ (n=32; median survival, 112.5 days) *vs* Tyr-Cre^ERT2^ *Braf*^V600E/+^ *Trp53*^fl/fl^ *Prex2*^-/-^ (n=17; median survival, 116 days) mice, *p*=0.4488. Bottom right panel, overall survival of Tyr-Cre^ERT2^ *Braf*^V600E/+^ *Pten*^fl/+^ *Prex2*^+/+^ (n=12; median survival, 65 days) *vs* Tyr-Cre^ERT2^ *Braf*^V600E/+^ *Pten*^fl/+^ *Prex2*^gd/gd^ (n=13; median survival, 64 days) mice, *p*=0.7544. *p*-values calculated by log-rank (Mantel–Cox) test.

Our search for new therapeutic approaches in melanoma has focussed upon the RAC1-GTPase signalling pathway. Integral to the activation of RAC1 signalling are the guanine nucleotide exchange factors (GEFs), such as the phosphatidylinositol-3,4,5-trisphosphate (PIP_3_)–dependent RAC-exchangers PREX1 and PREX2, which promote the conversion of RAC1 from an inactive GDP-bound form to an active GTP-bound conformation^10,11^. This switch potentiates binding of downstream effectors, such as the catalytic PI3K-isoform PI3Kβ/p110β, which dictate the output of RAC1-driven signalling^12^. We have demonstrated that the RAC1-GEF PREX1 plays critical roles in melanoblast migration during early murine embryonic development, melanoma cell invasion and migration, and metastasis in a patient-relevant preclinical model of *NRAS*-mutant melanoma^13^. Similarly, we demonstrated that RAC1 is required for embryonic melanoblast migration and function, melanoma growth *in vivo*, and tumour-cell invasion and migration^14,15^. (Fig 1A).

Although *PREX1* overexpression in melanoma can elicit invasion and metastasis, *PREX1* is rarely mutated in human cancer^13,16^. However, *bona fide* tumour-associated activating mutations in *RAC1* have been identified in human melanoma^17,18^, with the *RAC1*P29S hotspot mutation associated with resistance to BRAF-targeted therapies^19,20^. In addition, rearrangements, amplifications, missense and nonsense/truncating mutations in the RAC-GEF *PREX2* (a homolog of *PREX1*) are frequently detected in melanoma^21^. Nonetheless, no common or hotspot mutations in *PREX2* have been identified, and although some studies have indicated that truncating mutations may drive constitutive RAC-GEF activity^22,23^, the impact of *PREX2* mutation in melanoma remains contentious.

The tumour suppressor PTEN functions as a lipid phosphatase that dephosphorylates PIP_3_ to generate phosphatidylinositol-4,5-diphosphate (PIP_2_), thereby antagonising PREX2 activity^24^ and PI3K/AKT signalling. PTEN can also directly inhibit the GEF activity of PREX2 towards RAC1, suppressing cell migration and invasion^24^. The corollary of these findings is that *PTEN* loss would upregulate the GEF activity of PREX2 and, consequently, stimulate RAC1 function, rendering targeting of PREX2 in the context of PTEN-deficient melanoma a rational approach. Intriguingly, the aforementioned melanoma-associated RAC1-activating mutations appear to be mutually exclusive with *PTEN* loss, suggesting a scenario whereby loss of *PTEN* relieves the inhibition of PREX2^24,25^. Conversely, PREX2 can inhibit PTEN, promoting cell proliferation and tumorigenesis by activating downstream PI3K/AKT signalling^23,25,26^ (Fig. 1A).

Building on these observations, we use genetically engineered mouse models (GEMMs) and xenografts to characterise the role of PREX2 in BRAFV600E*-*driven melanoma *in vivo*. We further dissect the association between PREX2 and common tumour-suppressor gene mutations, such as *Trp53* or *Pten* loss and, perhaps more importantly, assess how deletion or inactivating mutation of *Prex2* impacts the therapeutic efficacy of MAPK-targeting agents in a preclinical setting. Our data suggest that while *Prex2* deletion has little impact upon tumour initiation, and progression, irrespective of tumour-suppressor gene mutation, it selectively sensitizes to MEK1/2 inhibition in the context of *Pten* deficiency. Moreover, our studies in GEMMs and human melanoma-derived cell lines suggest that co-targeting of the MAPK and RAC1/p110β signalling axes may be an efficacious therapeutic strategy in BRAFV600E-driven, PTEN-deficient melanoma.

## Results and discussion

### Deletion of *Prex2* does not impact development and progression of BRAFV600E-driven melanoma *in vivo*

While *PREX2* mutations are detected in ∼26% of human melanoma samples (Fig. 1B), these broadly lack functional annotation. Indeed, while *PREX2* is most frequently mutated in melanoma, distinct pan-cancer studies have identified widespread mutations across multiple tumour types^27^ (Supplementary Fig. S1A, B), although the biological and clinical significance of these mutations remains unclear. Biochemical analyses suggest that truncating *PREX2* mutations and a subset of missense mutations may have a pro-tumorigenic role^22,23^. This notion is confounded by clinical data from the TCGA PanCancer dataset, which suggest that the incidence of *PREX2* mutation is associated with extended progression-free survival in all cutaneous melanoma, irrespective of *BRAF* status (Fig. 1C, Supplementary Fig. S1C), but with no improvement in overall survival (Supplementary Fig. S1D, E). Notably, total mutation burden (TMB) appears to be elevated in *PREX2* mutant tumours and TMB is associated with improved overall survival (Supplementary Fig. S1F), suggesting that PREX2 mutation may be associated with a favourably prognostic mutator phenotype. Considering these observations, we sought to determine the functional role of PREX2 in melanoma *in vivo*, combining GEMMs of *BRAF*V600E-mutant melanoma with constitutive *Prex2* deletion. Transgenic expression of oncogenic *Braf*^V600E^ and/or deletion of tumour suppressor genes was targeted to the adult melanocyte population with a tamoxifen-inducible Cre-recombinase, under the control of the tyrosinase promoter (Tyr-Cre^ERT2^)^28^ (Fig. 1D, lower panels). The phenotype driven by these genetic aberrations in the melanocyte population has been well characterised in the mouse^5^, with the classical disease trajectory well understood (Fig. 1D, upper panels).

Melanocyte-specific expression of the BRAFV600E oncoprotein (Tyr-Cre^ERT2^ *Braf*^V600E/+^–henceforth BRAF) resulted in robust development of naevi 2–6 weeks post induction (median onset, 24 days), with lowly penetrant melanoma observed in ∼40% (9/22) of mice within 1 year (median onset, 403 days) (Supplementary Fig. S2A). Melanoma-bearing animals were aged to a defined endpoint (tumour diameter ≤ 15 mm, or ulceration), with an overall median survival of 466 days (Fig. 1E). To examine the role of PREX2 in melanoma, we interbred BRAF mice with a mouse line carrying germline deletion of *Prex2* (Tyr-Cre^ERT2^ *Braf*^V600E/+^ *Prex2*^-/-^ – henceforth BRAF PREX2). Compared with BRAF mice (median onset, 24 days), we observed a significant delay in naevus formation in BRAF PREX2 mice (median onset, 37 days), with no significant impact on tumour initiation (median tumour onset, 301 days *vs* 407 days), or overall survival (median survival, 375 days *versus* 466 days) (Fig. 1E, Supplementary Fig. S2A).

To generate more rapid and robust models of melanoma, we combined *Braf* mutation with targeted deletion of the key tumour suppressor genes *Pten* or *Trp53* – both common genetic events in the pathogenesis of melanoma in patients (Fig. 1A). Given previous reports of the reciprocal inhibition between PREX2 and PTEN^24,25^, and the loss of *PTEN* expression frequently observed in human melanoma^29^, we investigated the impact of *Prex2* deletion in models driven by loss of *Pten*, compared with loss of *Trp53* (Fig. 1E; Supplementary Fig. S2C, E). Alongside BRAFV600E, deletion of *Pten* (Tyr-Cre^ERT2^ *Braf*^V600E/+^ *Pten*^fl/+^– henceforth BRAF PTEN) or *Trp53* (Tyr-Cre^ERT2^ *Braf*^V600E/+^ *Trp53*^fl/fl^ – henceforth BRAF P53) did not significantly impact naevus formation (median onset – 32 and 28 days, respectively, compared to 24 days in BRAF cohort). However, deletion of either accelerated primary tumour development (median tumour onset – BRAF PTEN, 51 days; BRAF P53, 79 days; BRAF, 407 days) (Supplementary Fig. S2A, C, E), which translated into a significant decrease in median overall survival (BRAF PTEN, 81 days; BRAF P53, 113 days; BRAF, 466 days) (Fig. 1E). Deletion of *Prex2* did not impact the onset of naevi, primary tumour formation, or overall survival in either BRAF PTEN or BRAF P53 mice (Fig. 1E, Supplementary Fig. S2C, E). We observed suppressed PTEN expression in the tumour cells (but not stroma) of BRAF PTEN melanoma (and was unaffected by PREX2 loss), and nuclear accumulation of p21 was reduced in BRAF P53 melanoma (Supplementary Fig. S2B,D,F). PREX2 loss did not appear to markedly influence expression of these biomarkers in any model tested.

Our data indicate that PREX2 function is dispensable for early initiation and progression of BRAF-driven melanoma and do not support the existence of a melanomagenesis-relevant mutual inhibition between PTEN and PREX2 *in vivo*^24,25^.

### Loss-of-function mutation of *PREX2* phenocopies genetic deletion in BRAF PTEN melanoma

While PREX2 primarily functions as a GEF for the small GTPase RAC1, it also sits at the nexus of multiple complex signalling networks, it is conceivable that *Prex2* deletion may elicit off-target phenotypes. Therefore, we generated an enzymatically inactive, “GEF-dead” mutant PREX2 allele to determine whether functional loss phenocopies *Prex2* deletion *in vivo* (Supplementary Fig. 3). We combined this mutant *Prex2*^E22A,N204A^ (*Prex2*^gd^) allele with our BRAF PTEN melanoma model (Tyr-Cre^ERT2^ *Braf*^V600E/+^ *Pten*^fl/+^ *Prex2*^gd/gd^ – henceforth BRAF PTEN PREX2-GD) (Supplementary Fig. S2G). As observed in PREX2-deficient animals, this loss-of-function mutation had no impact upon the disease trajectory, with naevus onset, melanoma initiation, and overall survival of BRAF PTEN PREX2-GD animals equivalent to that of a BRAF PTEN control cohort (Fig. 1E, Supplementary Fig. S2G). Notably, the acceleration of naevus formation observed in BRAF PTEN PREX2 mice (Supplementary Fig. S2C) was not recapitulated in BRAF PTEN PREX2-GD cohorts (Supplementary Fig. S2G). Given that the PREX2-GD protein solely lacks GEF activity, this may be driven by loss of a non-GEF PREX2 function, such as the reported reciprocal inhibitory interaction between PREX2 and PTEN. This is supported by the observation that while *Prex2* deletion accelerated naevus onset in the context of heterozygous *Pten* deletion, it had no impact upon naevus formation following *Trp53* deletion (Supplementary Fig. S2C, E).

### PREX2 mutation or loss influences efficacy of MEK inhibition in BRAF-driven melanoma

Intriguingly, both loss-of-function mutation of PTEN, a suppressor of PREX2 activity, and activating mutation of RAC1, a downstream effector of PREX2, are known to modulate the response to BRAF inhibition^4,30,31^. We therefore addressed whether PREX2 impacts targeted therapeutic responses in melanoma. Given that MAPK-pathway activation is the key downstream effector of oncogenic BRAFV600E, and that effective targeting of this pathway is a current standard-of-care for *BRAF*-mutant melanoma, we tested the efficacy of a clinically relevant MEK1/2 inhibitor, selumetinib (AZD6244)^32^ (Fig. 1A), in PREX2-deficient *versus* -proficient BRAF PTEN and BRAF P53 melanoma. Mice were enrolled onto treatment having developed a single melanoma of diameter 7–10 mm (Supplementary Fig. S4B–D), and treated to clinical endpoint. AZD6244 had significant efficacy in BRAF PTEN mice, while in BRAF P53, despite short-term suppression of tumour growth, rapid regrowth suggested melanomas were either intrinsically resistant to treatment or capable of rapid reactivation of suppressed signalling pathways (Fig. 2A, B).

**Fig. 2:**
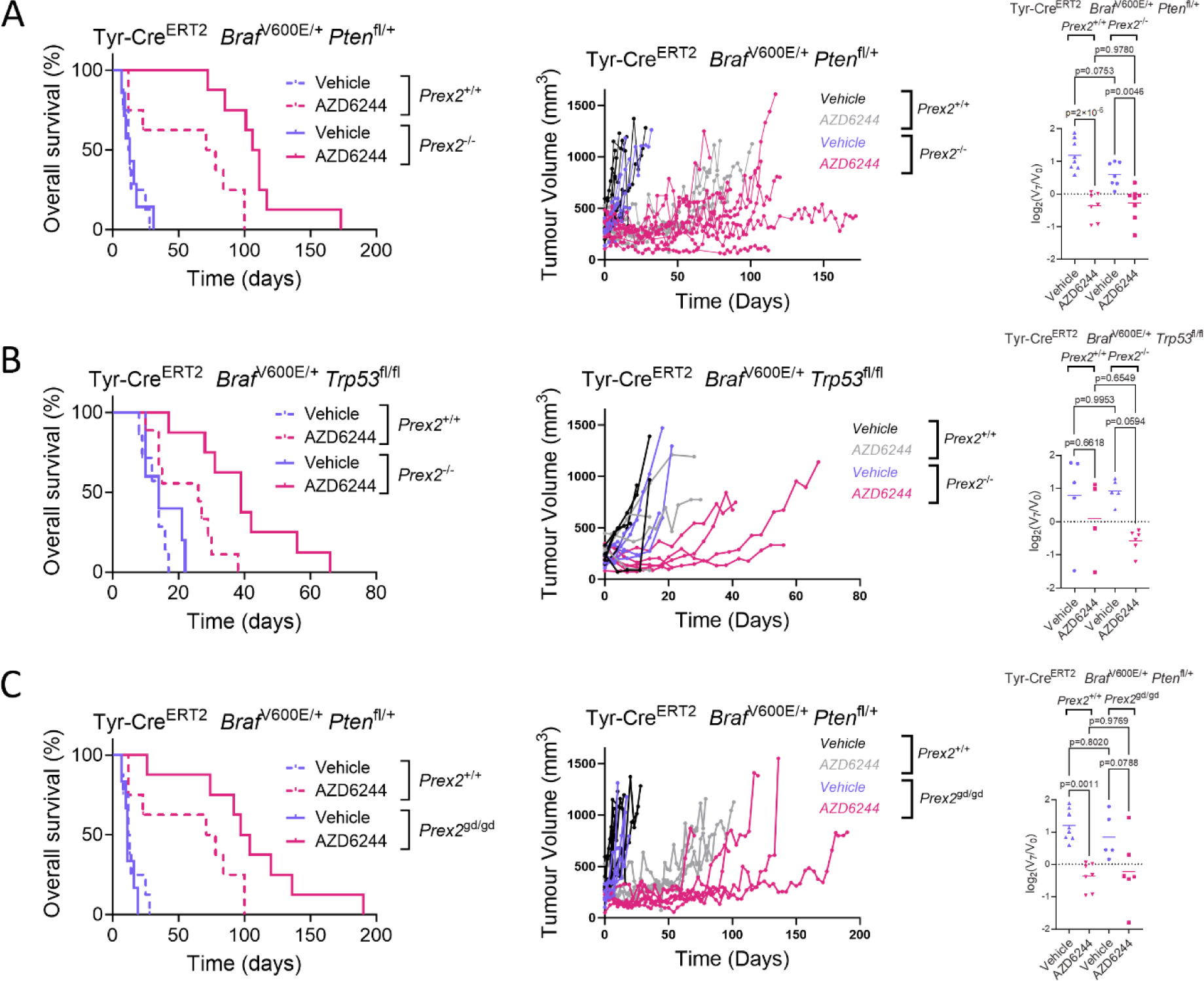
Loss of PREX2 function sensitises to MAPK inhibition. A) Left panel, Kaplan–Meier overall survival of mice with the indicated genotypes and treatments. BRAF PTEN+vehicle (n=8; median survival, 12.5 days) *vs* BRAF PTEN+AZD6244 (n=8; median survival, 74.5 days), p=0.0096; BRAF PTEN PREX2+vehicle, (n=7; median survival, 13 days) *vs* BRAF PTEN PREX2+AZD6244 (n=8; median survival, 108.5 days), p=0.00005; BRAF PTEN+AZD6244 (n=8; median survival, 74.5 days) *vs* BRAF PTEN PREX2+AZD6244 (n=8; median survival, 108.5 days), p=0.0021 Log-rank (Mantel–Cox) test. Centre panel, longitudinal growth of individual tumours from vehicle-treated (n=8) *vs* AZD6244-treated (n=7) BRAF PTEN cohorts and vehicle-treated (n=7) *vs* AZD6244-treated (n=8) BRAF PTEN PREX2 cohorts. Right panel, relative change in tumour volume in mice with the indicated genotypes over the first 7 days of indicated treatment. BRAF PTEN+vehicle (n=8), BRAF PTEN+AZD6244 (n=7), BRAF PTEN PREX2+vehicle (n=7), BRAF PTEN PREX2+AZD6244 (n=8). p-values were calculated by one-way ANOVA corrected for multiple comparisons (Tukey). B) Left panel, Kaplan–Meier overall survival of mice with the indicated genotypes and treatments. BRAF P53+vehicle (n=7; median survival, 14 days) *vs* BRAF P53+AZD6244 (n=9; median survival, 26 days), p=0.0204; BRAF P53 PREX2+vehicle (n=5; median survival, 14 days) *vs* BRAF P53 PREX2+AZD6244 (n=8; median survival, 39 days), p=0.0010; BRAF P53+AZD6244 (n=8; median survival, 14.5 days) *vs* BRAF P53 PREX2+AZD6244 (n=10; median survival, 35 days), p=0.0034. p-values calculated by log-rank (Mantel–Cox) test. Centre panel, longitudinal growth of individual tumours from vehicle-treated (n=5) *vs* AZD6244-treated (n=4) BRAF P53 cohorts and vehicle-treated (n=7) *vs* AZD6244-treated (n=6) BRAF P53 PREX2 cohorts. Right panel, relative change in tumour volume over the first 7 days of indicated treatment. BRAF P53+vehicle (n=5), BRAF P53+AZD6244 (n=4), BRAF P53 PREX2+vehicle (n=5), BRAF P53 PREX2+AZD6244 (n=6); p-values, one-way ANOVA corrected for multiple comparisons (Tukey). C) Left panel, Kaplan–Meier overall survival of BRAF PTEN and BRAF PTEN PREX2-GD cohorts treated with vehicle or AZD6244. BRAF PTEN+vehicle (n=8; median survival, 12.5 days) *vs* BRAF PTEN+AZD6244 (n=8; median survival, 74.5 days), p=0.0096; BRAF PTEN PREX2-GD+vehicle (n=6; median survival, 11 days) *vs* BRAF PTEN PREX2-GD+AZD6244 (n=8; median survival, 100.5 days), p=0.0002; BRAF PTEN+AZD6244 (n=8; median survival, 74.5 days) *vs* BRAF PTEN PREX2-GD^+^AZD6244 (n=8; median survival, 100.5 days), p=0.0414. p-values calculated by log-rank (Mantel–Cox) test. Centre panel, longitudinal growth of individual tumours from vehicle-treated (n=8) *vs* AZD6244-treated (n=7) BRAF PTEN cohorts and vehicle-treated (n=6) *vs* AZD6244-treated (n=8) BRAF PTEN PREX2-GD cohorts. Right panel, relative change in tumour volume over the first 7 days of indicated treatment. BRAF PTEN+vehicle (n=8), BRAF PTEN+AZD6244 (n=7), BRAF PTEN PREX2-GD+vehicle (n=5), BRAF PTEN PREX2-GD+AZD6244 (n=6). p-values calculated by one-way ANOVA corrected for multiple comparisons (Tukey). Note that the same BRAF PTEN treatment datasets are represented in B and D.

We next determined whether deletion of *Prex2* impacts the baseline therapeutic efficacy of AZD6244 in our melanoma models. BRAF PTEN PREX2 and BRAF P53 PREX2 tumours exhibited significant sensitivity to AZD6244, with equivalent early responses resulting in substantial tumour regression in both models, albeit with more prolonged sensitivity observed in BRAF PTEN PREX2 tumours (Fig. 2A, B, Supplementary Fig. S4A, B). These data represent the first indication that PREX2 may modify the response to MAPK-targeting therapies in BRAF-driven melanoma *in vivo*. We next assessed biomarker expression following short-term treatment with AZD6244. Here, mice were enrolled onto study, but sampled following 5 days of treatment. MAPK and mTORC1 signalling were suppressed in both the presence and absence of *Prex2*, indicated by reduced phosphorylation of ERK1/2 and ribosomal protein S6 (RPS6), at sites phosphorylated by MAPK and mTORC1 signalling respectively (Supplementary Fig. S4D).

PREX2 deficiency significantly extended the duration of response to MAPK-targeted therapy. The BRAF PTEN PREX2-GD model enabled us to test whether this phenotype was attributable to the RAC-GEF activity of PREX2. For this, we enrolled BRAF PTEN PREX2-GD mice onto AZD6244 or vehicle control, and assessed both the immediate and prolonged impact. Treatment of BRAF PTEN PREX2-GD mice with a MAPK inhibitor caused marked tumour regression, prolonged treatment response, and dramatically extended overall survival compared with vehicle control (Fig. 2C, Supplementary Fig. S4C). The extension of overall survival elicited by AZD6244-mediated MAPK-pathway inhibition, was similar in the BRAF PTEN PREX2 and BRAF PTEN PREX2-GD cohorts (95.5 days *vs* 89.5 days, respectively), and significantly greater than BRAF PTEN (62 days) (Fig. 2A, C). The observation that the PREX2-GD GEF-dead mutant phenocopies *PREX2* deletion suggests that the canonical GEF activity of PREX2 counteracts MAPK inhibition in BRAF PTEN melanoma. In turn, this implicates RAC1 signalling in the resistance to MAPK-targeted therapy and supports the reported role for activating *RAC1*P29S mutations in melanoma resistance to vemurafenib^31^.

### Genetic and pharmacological targeting of p110β phenocopies PREX2 mutation/loss in PTEN-deficient melanoma

Our data indicate that PTEN/PREX2/RAC1 signalling impacts response to MAPK-targeting therapies in melanoma, with the corollary that a combination of therapeutics targeting both MAPK and PTEN/PREX2/RAC1 signalling may be an effective rational approach. While the established standard-of-care for *BRAF*-mutant melanoma includes MAPK targeting, the direct therapeutic targeting of small GTPases, such as RAC1, has been viewed as challenging – despite the significant recent success of small molecules targeting oncogenic RAS^33^. We therefore sought to identify key alternative targetable signalling nodes associated with the PTEN/PREX2/RAC1 signalling axis that could be exploited for combination therapies with MAPK inhibitors. One such target of interest is the type I PI3K isoform p110β (encoded by *PIK3CB*). This not only acts as a downstream effector of RAC1^12^ but, akin to PREX2, is activated by the binding of the Gβγ subunit of G-protein–coupled receptors^34,35^ and exhibits enriched catalytic activity in PTEN-deficient tumours^36,37^.

The association between RAC1 and p110β, which stimulates RAC1 catalytic activity, is mediated by the canonical RAS binding domain (RBD) of p110β. We reasoned that if the interaction between RAC1 and p110β is important for modulating melanoma response to MAPK-targeting therapies, its genetic disruption might phenocopy *PREX2* loss/mutation and sensitize to therapy. To test this hypothesis, we interbred mice carrying the mutant *Pik3cb*^S205D,K224A^ allele (*Pik3cb*^rbd^), which encodes p110β with a non-functional RBD knocked-in to the endogenous *Pik3cb* locus^12^, with our BRAF PTEN melanoma model, generating Tyr*-*Cre^ERT2^ *Braf*^V600E/+^ *Pten*^fl/+^ *Pik3cb*^rbd/rbd^ mice (henceforth BRAF PTEN PIK3CBmut) (Fig. 3A, Supplementary Fig. S5A). This loss-of-function *Pik3cb* mutant had little-to-no impact upon disease trajectory, with naevus onset and melanoma initiation similar to BRAF PTEN controls, although *Pik3cb* mutation did appear to result in improved overall survival (Supplementary Fig. S5A). In line with the prediction that disrupting the RAC1–p110β interaction might phenocopy deletion or inactivating mutation of *Prex2*, melanomas arising in the BRAF PTEN PIK3CBmut model were acutely sensitive to MAPK inhibition in the short term, and mice exhibited prolonged overall survival and a delay in the onset of therapeutic resistance compared with BRAF PTEN controls (Fig. 3A, Supplementary Fig. S5B).

**Fig. 3:**
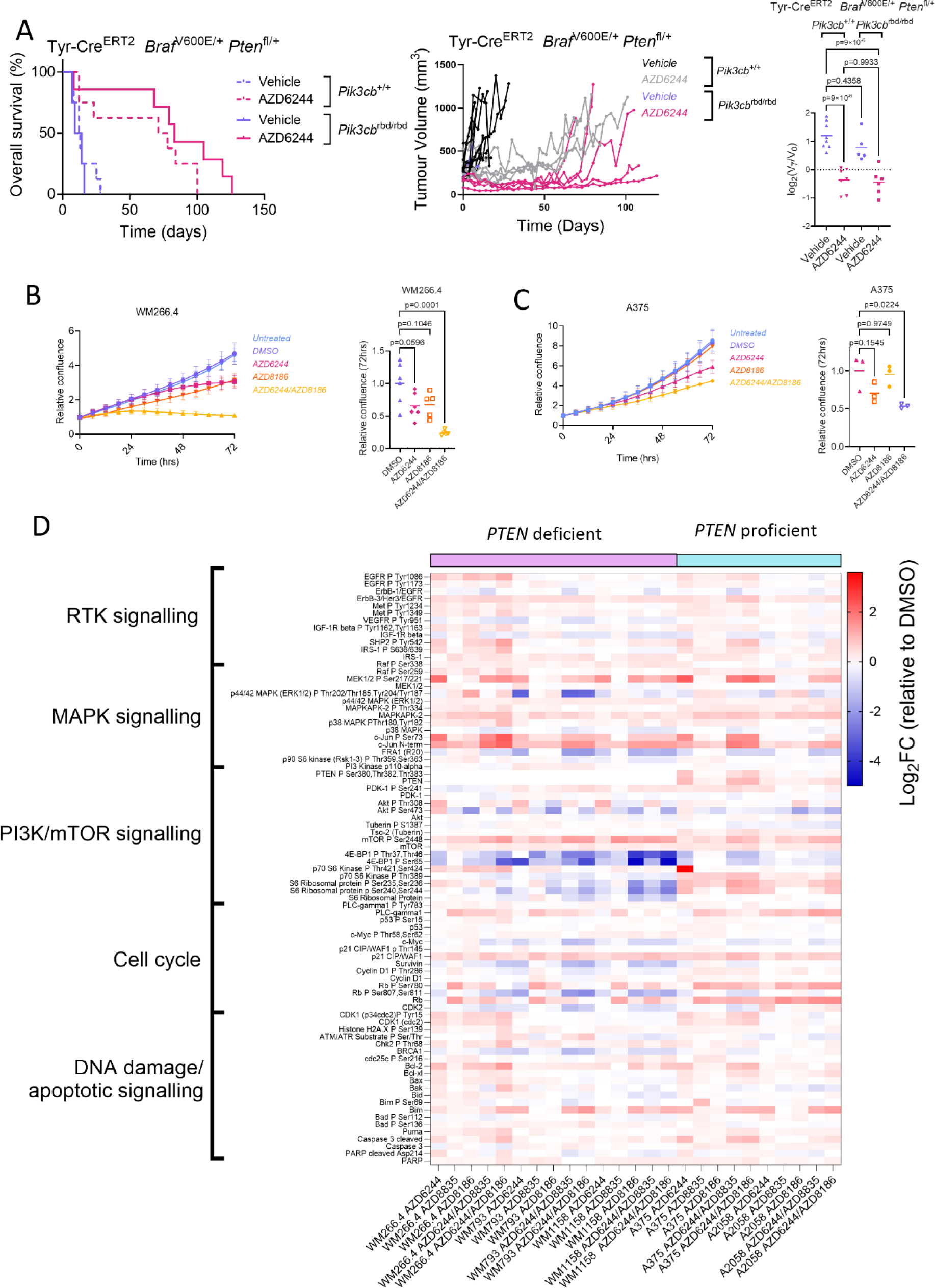
Functional loss of PIK3CB sensitises to MAPK inhibition and suppresses mTOR signalling. A) Left panel, Kaplan–Meier overall survival of BRAF PTEN mice treated with vehicle or AZD6244. BRAF PTEN+vehicle (n=8; median survival, 12.5 days) *vs* BRAF PTEN+AZD6244 (n=8; median survival, 74.5 days), p=0.0096; BRAF PTEN PIK3CB-mut+vehicle (n=4; median survival, 11.5 days) *vs* BRAF PTEN PIK3CB-mut+AZD6244 (n=7; median survival, 83 days), p=0.0082; BRAF PTEN+AZD6244 (n=8; median survival, 74.5 days) *vs* BRAF PTEN PIK3CB-mut+AZD6244 (n=7; median survival, 83 days), p=0.1643. p-values calculated by log-rank (Mantel–Cox) test. Centre panel, longitudinal growth of individual tumours from vehicle-treated (n=8) *vs* AZD6244-treated (n=7) BRAF PTEN cohorts and vehicle-treated (n=5) *vs* AZD6244-treated (n=7) BRAF PTEN PIK3CB-mut cohorts. Right panel, relative change in tumour volume over the first 7 days of indicated treatment. BRAF PTEN+vehicle (n=8), BRAF PTEN+AZD6244 (n=7), BRAF PTEN PIK3CB-mut+vehicle (n=5), BRAF PTEN PIK3CB-mut+AZD6244 (n=6); p-values calculated by one-way ANOVA corrected for multiple comparisons (Tukey). Centre line, mean. Note that the BRAF PTEN treatment datasets are reproduced from Fig. 2. B) Left panel, relative confluence of WM266.4 cells treated with indicated treatments over time. Representative of a minimum of 5 independent experiments and 3 technical replicates. Data, mean ± SEM (confluence relative to starting point). Right panel, relative change in confluence of WM266.4 cells over indicated 72 h treatment. Centre line, mean. p-values calculated by one-way ANOVA corrected for multiple comparisons (Tukey). C) Left panel, relative confluence of A375 cells treated with indicated treatments over time. Representative of 3 independent experiments with a minimum of 3 technical replicates. Data, mean ± SEM (confluence relative to starting point). Right panel, relative change in confluence of A375 cells over indicated 72 h treatment. Centre line, mean. p-values calculated by one-way ANOVA corrected for multiple comparisons (Tukey). D) RPPA dataset comparing PTEN-deficient and PTEN-proficient melanoma cell lines treated with indicated treatments. Cell line names and treatments are indicated below the heatmap. Antigens, detected by RPPA antibodies, are listed vertically according to biological process/signalling pathway. Colour intensity scale indicates high (red) and low (blue) log_2_FC of RPPA intensity values relative to the relevant DMSO control. Results are representative of 2 technical and 5 biological replicates per condition.

### Combined MEK and p110β inhibition suppresses both proliferation and mTORC1 activity in human PTEN-deficient melanoma cells *in vitro*

Given the tumour regression observed in all AZD6244-treated BRAF PTEN melanoma models and the prolonged sensitivity to MAPK inhibition, driven by genetic targeting of *Prex2* or *Pik3cb*, we interrogated the underlying mechanisms in established human melanoma cells. We compared the *in vitro* growth kinetics of a *BRAF*-mutant, *PTEN*-deficient melanoma cell line (WM266.4) and a *BRAF*-mutant, *TP53*-deficient but *PTEN*-proficient line (A375) following treatment with AZD8186, a clinically relevant, isoform-specific inhibitor of p110β/δ^38^. We used IncuCyte timelapse microscopy to measure real-time changes in the relative confluence of cell cultures treated with vehicle or drug. Cells were seeded 24 h prior to treatment, with confluence measurements subsequently performed at 6-h intervals. Combined targeting of MEK1/2 and p110β/δ with AZD6244 and AZD8186 almost completely abrogated the growth of PTEN-deficient WM266.4 cells over the same period, in contrast to treatment with AZD6244 alone (Fig. 3B, C). This combination treatment also elicited a slight, yet significant, suppression of growth of the PTEN-proficient line A375 (Fig. 3C). Subsequent experiments demonstrated that this effect was selectively for inhibition of p110β over p110α, could be recapitulated by inhibition of mTOR kinase with AZD2014/vistusertib, and was not observed in the PTEN-deficient melanoma line WM793 (Supplementary Fig. S5C-E).

This confirmed that the treatment of established human melanoma-derived cell lines *in vitro* recapitulates responses *in vivo*, providing a tractable platform for interrogating the mechanistic implications of co-targeting MAPK and PREX2/RAC1/p110β. We next performed a targeted proteomics approach via reverse-phase protein array (RPPA), allowing assessment of key signalling nodes across a panel of PTEN-deficient and - proficient lines upon treatment *in vitro*. Here, we used a broader collection of established melanoma lines, again encompassing PTEN-deficient (WM266.4, WM793, WM1158) and PTEN-proficient (A375, A2058) cells, and sought to assess PI3K isoform selectivity through head-to-head comparison of anti-proliferative and pro-apoptotic efficacy of the p110α-specific inhibitor AZD8835, with AZD8186, using the same experimental approach.

We assessed treatment impact across all lines on >50 key signalling nodes, subdivided into 5 broad classes – receptor tyrosine kinase signalling, MAPK signalling, PI3K/mTOR signalling, cell-cycle control, and DNA damage/apoptotic signalling. This allowed us to identify key differentiators of response between PTEN-deficient and - proficient lines as well as mediators of the selective response to p110β/δ inhibition over p110α inhibition. We observed differential responses to PI3K targeting in PTEN-proficient *versus* -deficient lines, whereby combination of either PI3K-targeting agent with AZD6244 had no additional impact upon any individual target or target class than treatment with AZD6244 alone. This suggests that activation of PI3K signalling does not represent a critical molecular response to MAPK inhibition in PTEN-proficient lines *in vitro* (Fig. 3D). As a counterpoint to these findings, we observed potentiation of AKT phosphorylation at the canonical PDK1 (Thr308) and mTORC2 (Ser473) target sites in AZD6244-treated PTEN-deficient (but not PTEN-proficient) lines, suggesting activation of PI3K signalling in response to MAPK inhibition in a *PTEN*-dependent manner. Notably, in all PTEN-deficient lines tested, p110β/δ inhibition with AZD8186 was more effective at suppressing AZD6244-mediated AKT phosphorylation at either site than p110α inhibition, suggesting that p110β/δ plays a dominant role in the activation of PI3K/AKT signalling in a PTEN-deficient setting. Our RPPA analyses also showed that inhibition of MEK1/2 and p110β/δ, through co-administration of AZD6244 and AZD8186, impacted several cellular signalling pathways/networks by suppressing key nodes controlling PI3K/mTOR signalling and cell-cycle progression. Amongst these, suppression of downstream targets/effectors of mTORC1, such as phosphorylation of 4EBP1 (Thr37/46 and Ser65) or RPS6 (Ser235/236 and Ser240/244), and suppression of key cell-cycle control nodes, such as expression of c-MYC and survivin, or phosphorylation of cyclin D1 (Thr286) and Rb (Ser807/811), suggested that p110β/δ inhibition may counteract AKT/mTORC1-dependent cell-cycle progression potentiated by MAPK inhibition (Fig. 3D).

We confirmed biomarker responses in pathways downstream of MAPK and PI3K signalling and validated our RPPA analysis *via* immunoblotting of protein lysates from vehicle- and drug-treated PTEN-deficient WM266.4 and WM793 lines and PTEN-proficient A375 cells. Biomarker responses were also compared at key signalling nodes in the PI3K/mTOR pathway in response to AZD6244-mediated MAPK inhibition combined with inhibitors of p110α (AZD8835), p110β/δ (AZD8186), AKT (AZD5363/capivasertib), or mTOR kinase (AZD2014/vistusertib) (Fig. 4A, Supplementary Fig. S6A–C). This demonstrated that AKT (Ser473) phosphorylation was induced in response to treatment with AZD6244, only in PTEN-deficient lines, with this phosphorylation sensitive to AZD8186 (Fig. 4A) and AZD2014 (Supplementary Fig. S6C), implicating p110β/δ and mTORC2, but not p110α in AKT activation in this setting. Similarly, RPS6 phosphorylation at key sites (Ser235/236 and Ser240/244) was most markedly suppressed in WM266.4 cells in response to combined treatment with AZD6244 and AZD8186 (Fig. 4A), recapitulating the RPPA analysis (Fig. 3D), but was also responsive to combined MEK/AKT and MEK/mTOR but not MEK/p110α inhibition (Supplementary Fig. S6A–C), suggestive of p110β/δ–AKT–mTORC1-dependent regulation.

**Fig. 4:**
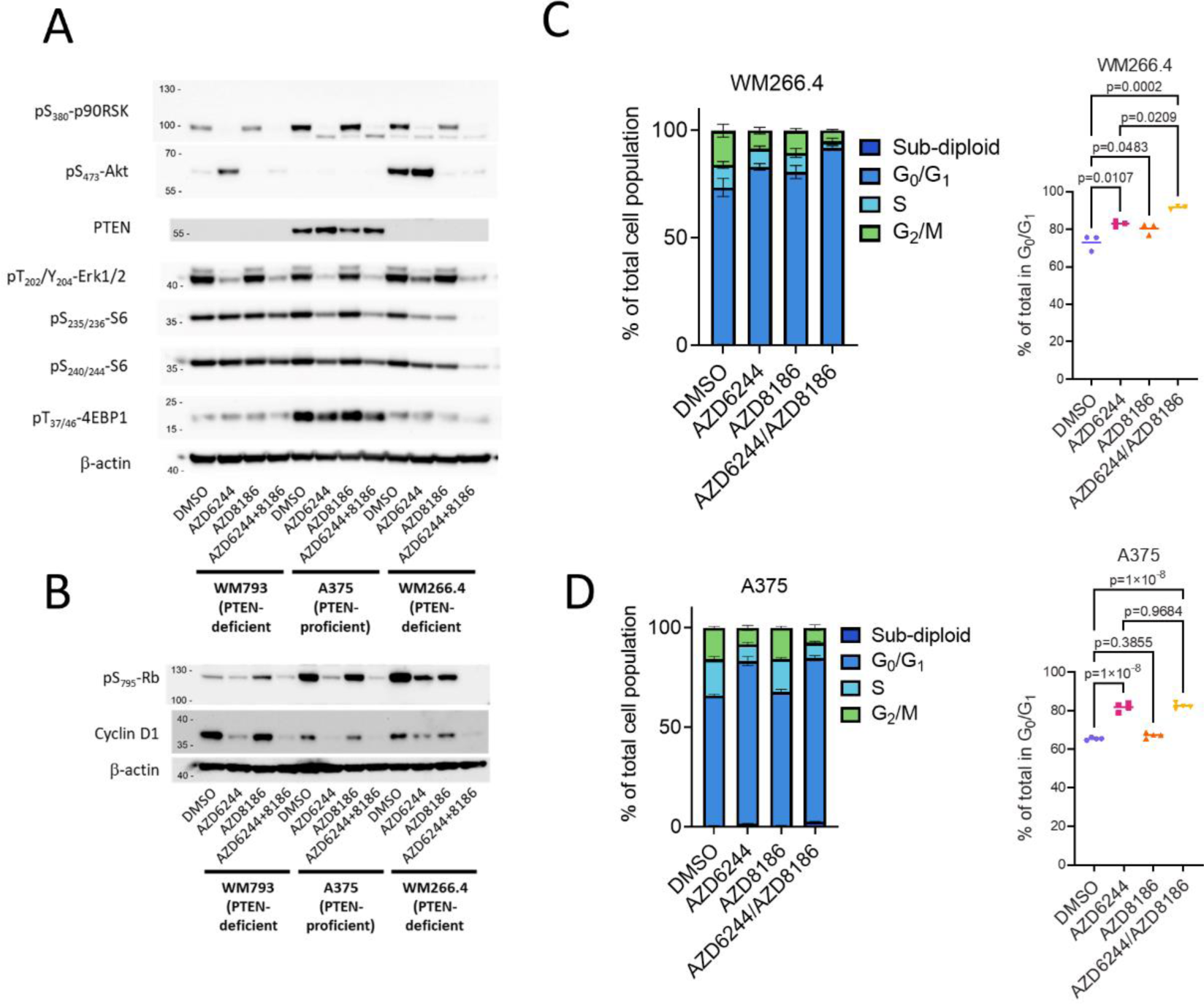
Co-targeting of MEK1/2 and p110β suppresses cell-cycle progression in human melanoma cells. A) Immunoblotting for indicated activated components of the MAPK–PI3K–mTOR pathway in human melanoma cells treated with vehicle or the indicated targeted therapeutics. β-actin serves as a sample integrity control. The blots are representative of 3 repeated experiments. B) Immunoblotting for activation of the CDK4/6–cyclin D1–Rb pathway in human melanoma cells treated with vehicle or the indicated targeted therapeutics. β-actin serves as a sample integrity control. The blots are representative of 3 repeated experiments. C) Left panel, Flow cytometry–based cell-cycle profiling of WM266.4 cells following indicated 24 h treatment. Right panel, proportion of WM266.4 cells in G_1_/S at 24 h. n = 3 independent experiments, p-values calculated by one-way ANOVA corrected for multiple comparisons (Tukey). D) Left panel, Flow cytometry–based cell-cycle profiling of A375 cells following indicated 24 h treatment. Right panel, proportion of A375 cells in G_1_/S at 24 h. n = 4 independent experiments, p-values calculated by one-way ANOVA corrected for multiple comparisons (Tukey).

Monotherapy with AZD6244 or AZD8186 elicited a predictable biomarker response, MEK1/2 inhibition suppressed phosphorylation of ERK1/2 and p90RSK phosphorylation but, in the absence of p110β/δ inhibition, this was not sufficient to curtail RPS6 phosphorylation (Fig. 4A). Indeed, in the PTEN-proficient A375 line, combined inhibition of MEK1/2 and p110α, p110β/δ, or AKT had very little additional impact on RPS6 phosphorylation, although the combination of the mTOR-kinase inhibitor AZD2014 with AZD6244 suppressed RPS6 phosphorylation even in this PTEN-proficient setting (Fig. 4A, Supplementary Fig. S6A–C). These data indicate substantial and complex crosstalk/compensation between the MAPK and PI3K/AKT/mTOR pathways in these cell lines and underscore the dependency of PTEN-deficient melanoma lines on p110β/δ signalling.

In addition to the suppression of PI3K/mTORC1 signalling, RPPA analysis indicated decreased cell-cycle progression (decreased phospho-Rb) following combined inhibition of MEK1/2 and p110β/δ selectively in PTEN-deficient lines (Fig. 3D). This was confirmed by immunoblotting, with the combination of AZD6244/AZD8186 resulting in ablation of Rb (Ser795) phosphorylation and, downregulation of cyclin D1 (Fig. 4B). Intriguingly, AZD6244-mediated inhibition of MAPK signalling suppressed Rb phosphorylation and cyclin D1 expression in PTEN-proficient A375 cells, but AZD8186 had no additive effect (Fig. 4B). Irrespective of *PTEN* status, MAPK inhibition attenuated Rb phosphorylation and cyclin D1 expression; notably, however, in PTEN-deficient WM266.4 cells, p110β/δ inhibition had an additive effect, further decreasing phospho-Rb and cyclin D1 levels (Fig. 4B). These observations suggest that MAPK and PI3Kβ signalling converge upon Rb *via* different mechanisms, such as transcriptional control of *CCND1* (encoding cyclin D1) by ERK1/2 and/or MYC, downstream of MAPK signalling, *versus* proteolytic degradation of cyclin D1 by GSK3β^39^, the kinase activity of which is in turn inhibited by PI3K/AKT/mTOR signalling^40^.

Our data further suggest that combined MEK/p110β inhibition may selectively and additively suppress the proliferation of WM266.4 cells (Figs. 3D and 4B) and, by extension, PTEN-deficient melanoma. Given the observed impact on mTORC1 signalling and Rb phosphorylation, it is likely that these therapies may also impact cell-cycle progression, which would account for the observed growth defect *in vitro*. Therefore, we next examined the impact of each MAPK and PI3K/AKT/mTOR mono- or combination therapy on cell-cycle progression through flow cytometry of synchronous WM266.4 (Fig. 4C, Supplementary Fig. S6D) or A375 (Fig. 4D, Supplementary Fig. S6E) cultures. MAPK inhibition increased the proportion of cells in G_0_/G_1,_ with a concomitant decrease in the number of cells in the S- and G_2_/M-phases, 24 h post treatment. Crucially, while the addition of AZD8186 had no further impact on any A375 subpopulation, it substantially increased the proportion of WM266.4 cells in G_0_/G_1_, compared with MAPK inhibition alone (Fig. 4C, D – right panels). Notably, MAPK inhibition was sufficient to drive an increase in the sub-diploid, apoptotic population in the A375 line, which was absent in WM266.4 cells (Fig. 4C, D – left panels). The impact on cell-cycle progression of adding AZD8186 (p110β/δ) to AZD6244 was phenocopied by with the addition of AZD2014 (mTOR) or AZD5363 (AKT), but not AZD8835 (p110α), in the PTEN-deficient WM266.4 line (Supplementary Fig. S6D – upper panels). Furthermore, combination treatment had no additional impact on the AZD6244-induced cell-cycle arrest of PTEN-proficient A375 cells (Supplementary Fig. S6D – lower panels). Notably, the PTEN-deficient WM793 line appeared resistant to cell-cycle arrest and apoptosis induction upon PI3K/AKT/mTOR targeting, with no greater efficacy of co-targeting than with AZD6244 monotherapy (Supplementary Fig. S6E-F), consistent with the lack of impact observed via confluence measurements (Supplementary Fig S5E). The resistance phenotype of WM793 cells could be explained, in part, by a mutation in *CDK4* (K22Q) predicted to uncouple mTORC1/cyclin-D1 from Rb phosphorylation and cell-cycle control, suggesting a mechanism of resistance to AZD6244/AZD8186 treatment in PTEN-deficient lines harbouring *CDK4* activating mutation/overexpression. These data are consistent with the observed pattern of Rb phosphorylation *in vitro*, the induction of a G0/G1 arrest following MAPK inhibition, and the selective anti-growth efficacy of combined MAPK and PI3Kβ inhibition in a subset of PTEN-deficient melanomas.

### Genetic ablation of *PREX1/2 in vitro* phenocopies genetic or pharmacological targeting of *Pik3cb*/p110β

While there is substantial evidence in the literature for a relationship between PREX2 and PTEN ^23–26^, and of regulatory/effector networks shared by PREX2 and p110β ^12,26,34,35,41^, it is not yet clear whether the phenotypes driven by PREX2 loss and p110β targeting are related. To address this question, we took advantage of CRISPR/Cas9-mediated gene editing to disrupt the expression of *PIK3CB*, *PREX2* or its close relative *PREX1* in the PTEN-deficient WM266.4 melanoma line and treated these CRISPR/Cas9-derived cell lines with AZD6244 and/or AZD8186. We hypothesised that *PIK3CB* or *PREX2* disruption would acutely sensitise WM266.4 lines to MEK inhibition with AZD6244 and, equally, would attenuate any survival benefit bestowed by AZD8186-mediated p110β/δ inhibition. We also chose to target *PREX1* as we have previously demonstrated that *PREX1* is highly expressed in WM266.4 cells^13^, and that it is both structurally and functionally homologous to PREX2 and may therefore compensate for PREX2 loss-of-function.

We generated polyclonal populations of CRISPR/Cas9-edited cells and verified successful editing by Sanger sequencing of the targeted region and the Tracking of Indels by Decomposition (TIDE) algorithm^42^. Following CRISPR/Cas9 genome editing, we used TIDE to calculate that 73.2% of *PIK3CB* transcripts, 87.1% (D2) or 64.3% (A4) of *PREX2* transcripts, and 81.6% (B4) or 82.6 (A4) of *PREX1* transcripts in the edited cell populations were generated from disrupted genomic sequences in their respective polyclonal cell line pools (Supplementary Fig. S7A–C).

We next used IncuCyte live-cell imaging to measure real-time changes in the relative cell confluence and growth kinetics of each edited cell line, treated with vehicle or drug, alongside the unedited parental WM266.4 cells. As expected, this analysis demonstrated that PIK3CB expression was dispensable for unperturbed WM266.4 growth, and that *PIK3CB* disruption phenocopied the p110β/δ inhibition induced by AZD6244-mediated MAPK inhibition (Fig. 5A, B). Crucially, *PIK3CB* disruption also abrogated any additional benefit provided by AZD8186, in combination with AZD6244, suggesting that, in this setting, the additive anti-proliferative impact of AZD8186 is mediated by the inhibition of p110β rather than p110δ (Fig. 5B). As with PIK3CB, PREX2 and PREX1 expression appeared dispensable for unperturbed growth of WM266.4 cells; indeed, as expected, we observed a significant potentiation of AZD6244-mediated growth inhibition of individually edited cells, albeit more modestly than with disruption of *PIK3CB* (Fig. 5B–D). Using the same gRNAs to disrupt *PREX2* and *PREX1*, we performed an independent round of CRISPR/Cas9 editing, with the resulting genome-edited polyclonal lines yielding a similarly modest potentiation of AZD6244 efficacy (Supplementary Fig. S7D, E).

**Fig. 5:**
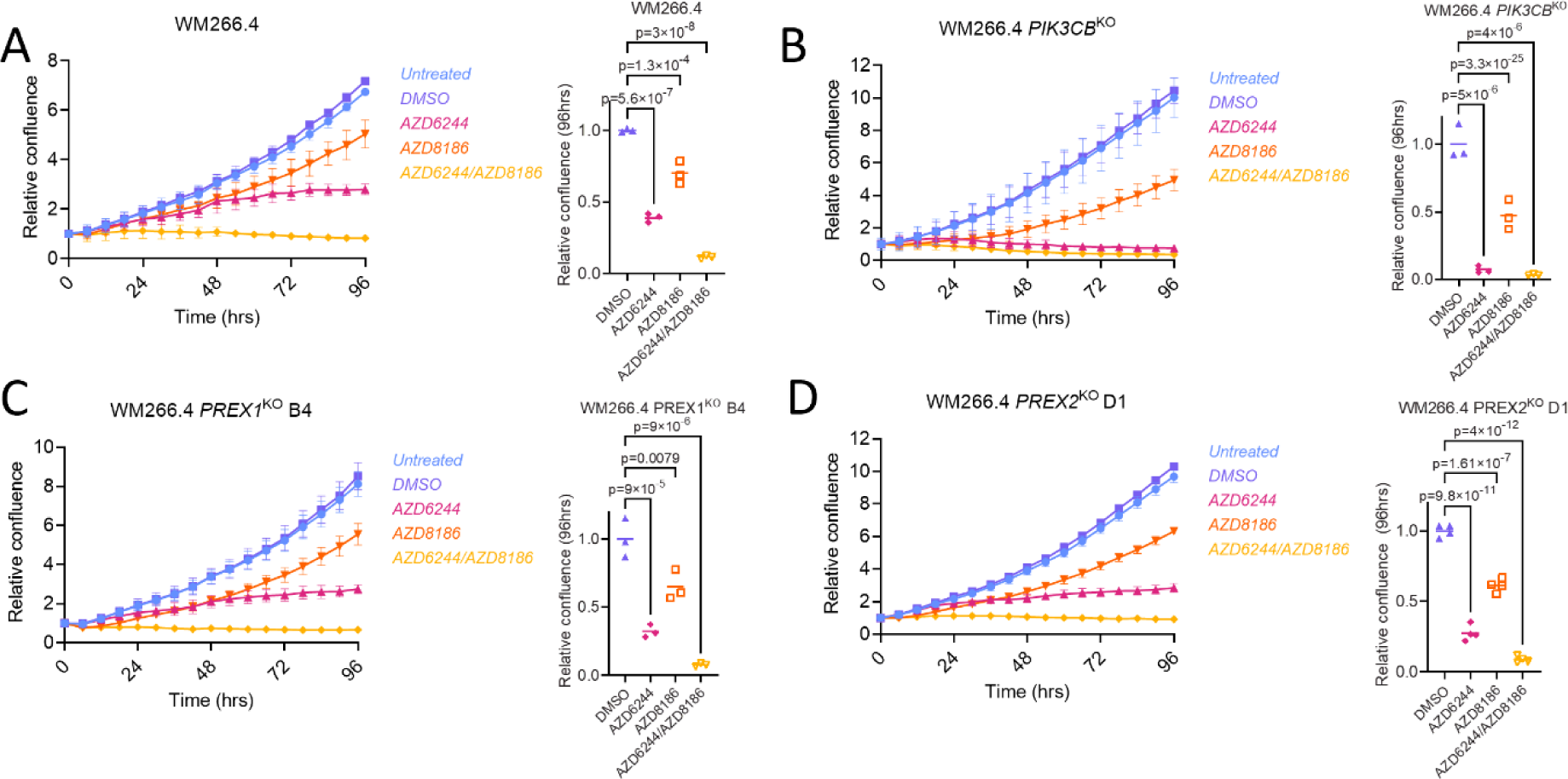
Genetic disruption of *PIK3CB*, *PREX1,* and/or *PREX2* drives sensitivity to MAPK inhibition *in vitro*. A) Left panel, relative confluence of parental WM266.4 cells treated with indicated treatments over time. Right panel, relative change in confluence of parental WM266.4 cells over indicated 96 h treatment. Centre line, mean. p-values calculated by one-way ANOVA corrected for multiple comparisons (Tukey). B) Left panel, relative confluence of WM266.4 *PIK3CB*^KO^ cells treated with indicated treatments over time. Right panel, relative change in confluence of WM266.4 *PIK3CB*^KO^ cells over indicated 96 h treatment. Centre line, mean. p-values calculated by one-way ANOVA corrected for multiple comparisons (Tukey). C) Left panel, relative confluence of WM266.4 *PREX1*^KO^ cells treated with indicated treatments over time. Right panel, relative change in confluence of WM266.4 *PREX1*^KO^ cells over indicated 96 h treatment. Centre line, mean. p-values calculated by one-way ANOVA corrected for multiple comparisons (Tukey). D) Left panel, relative confluence of WM266.4 *PREX2*^KO^ cells treated with indicated treatments over time. Right panel, relative change in confluence of WM266.4 *PREX2*^KO^ cells over indicated 96 h treatment. Centre line, mean. p-values calculated by one-way ANOVA corrected for multiple comparisons (Tukey). In all left panels, representative curves from 3 independent experiments are shown, with each datapoint representing the mean ± SEM of a minimum of 4 technical replicates of confluence relative to starting point. In all right panels, centre line represents the mean.

### Co-targeting of MEK1/2 and p110β has therapeutic efficacy in human PTEN-deficient melanoma xenografts *in vivo*

We next sought to test whether the identified association between *Prex2*/*Pik3cb* status and drug sensitivity is recapitulated *in vivo*. To do so, WM266.4 or A375 cells were engrafted subcutaneously into the flank of athymic CD1-*Foxn1^nu^* mice, with subsequent tumour outgrowth monitored over time. Mice were enrolled into appropriate treatment groups once engrafted tumours had reached a diameter > 7mm (Supplementary Fig. S8C–D), and therapeutic response was measured both in terms of primary tumour growth/regression and overall survival.

In contrast to our *in vitro* data, it was notable that MEK1/2 inhibition alone had an immediate impact on tumour growth in both models, markedly slowing the growth of WM266.4 xenografts and driving regression of A375 xenografts over the first 21 and 14 days of treatment, respectively (Fig. 6A, Supplementary Fig. S8A). Nonetheless, combined MEK1/2 and p110β inhibition had no additive effect on tumour volume, over MEK1/2 inhibition alone, in tumours arising from engrafted PTEN-proficient A375 cells and only modestly increased overall survival (Supplementary Fig. S8A). As predicted by our prior *in vitro* and *in vivo* data, however, the same inhibitor combination (AZD6244/AZD8186) demonstrated marked additional benefit, compared with MEK1/2 inhibition alone, driving tumour regression of PTEN-deficient WM266.4 xenografts and significantly increasing overall survival (Fig. 6A).

**Fig. 6:**
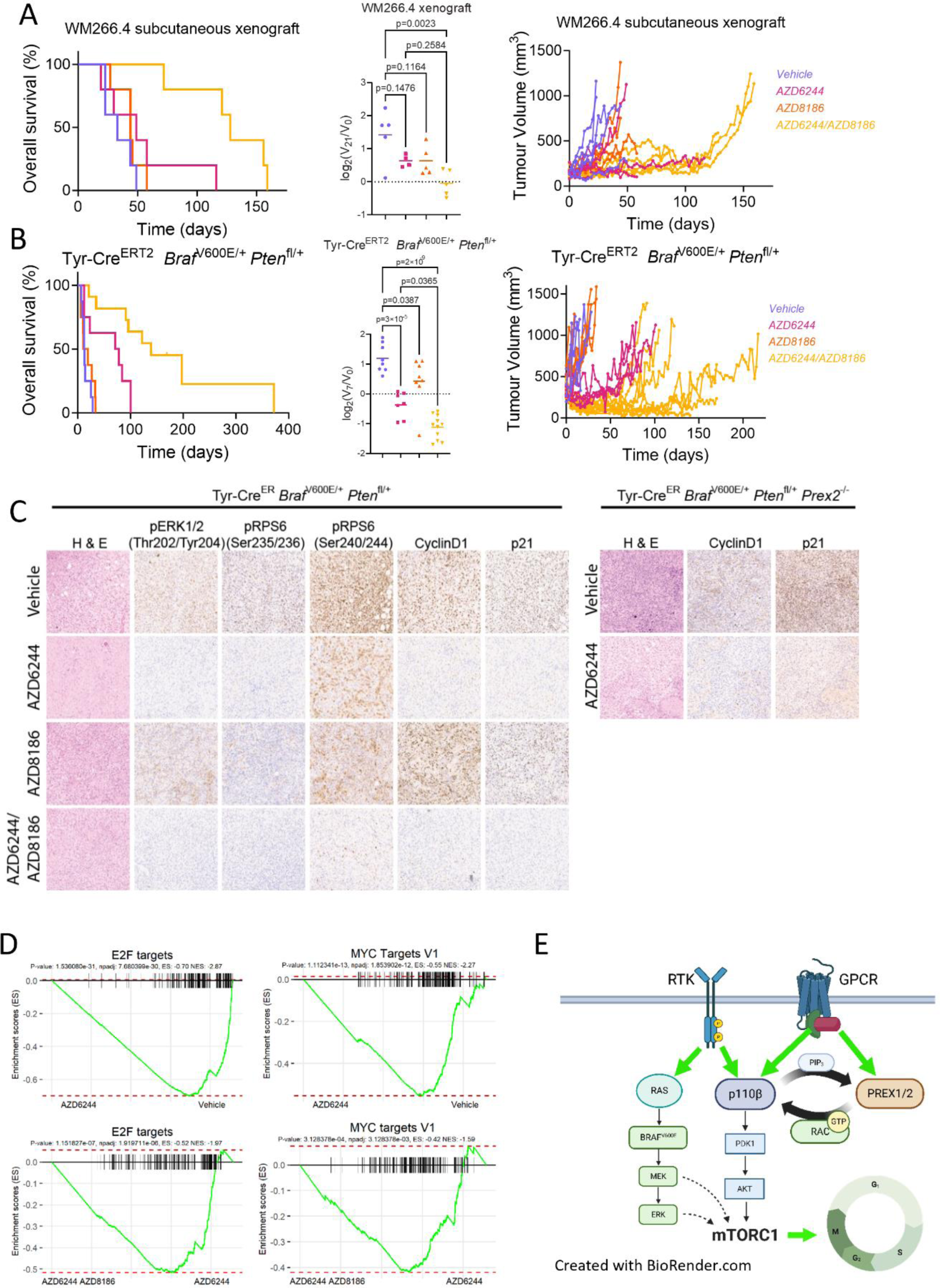
Co-targeting of MEK1/2 and p110β has therapeutic efficacy in melanoma *in vivo*. A) Left panel, Kaplan–Meier overall survival of mice harbouring WM266.4 subcutaneous xenografts treated with vehicle (n=5; median survival, 33 days), AZD6244 (n=5; median survival, 49 days), AZD8186 (n=5; median survival, 44 days), or AZD6244/AZD8186 (n=5; median survival, 128 days). p-values calculated by log-rank (Mantel–Cox) test – vehicle vs AZD6244, p=0.2269; vehicle vs AZD8186, p=0.3091; vehicle vs AZD6244/AZD8186, p=0.0017 and AZD6244 vs AZD6244/AZD8186, p=0.0064. Centre panel, relative change in tumour volume of WM266.4 xenografts over the first 21 days of indicated treatment. Vehicle (n=5), AZD6244 (n=4), AZD8186 (n=5), AZD6244/AZD8186 (n=5), p-values calculated by one-way ANOVA corrected for multiple comparisons (Tukey). Right panel, longitudinal growth of individual WM266.4 xenografts treated with vehicle (n=5) *vs* AZD6244 (n=5), AZD8186 (n=5), or AZD6244/AZD8186 (n=5). B) Left panel, Kaplan–Meier overall survival of BRAF PTEN mice treated with vehicle (n=8; median survival, 12.5 days), AZD6244 (n=8; median survival, 74.5 days), AZD8186 (n=8; median survival, 15 days), or AZD6244/AZD8186 (n=8; median survival, 139 days). p-values calculated by log-rank (Mantel–Cox) test – vehicle vs AZD6244, p=0.009550; vehicle vs AZD8186, p=0.437704; vehicle vs AZD6244/AZD8186, p=0.000012 and AZD6244 vs AZD6244/AZD8186, p=0.004813. Centre panel, relative change in tumour volume of BRAF PTEN mice over the first 7 days of indicated treatment. Vehicle (n=8), AZD6244 (n=7), AZD8186 (n=8), AZD6244/AZD8186 (n=11); p-values calculated by one-way ANOVA corrected for multiple comparisons (Tukey). Right panel, longitudinal growth of individual tumours from vehicle-treated (n=8) *vs* AZD6244-treated (n=7), AZD8186-treated (n=8), and AZD6244/AZD8186-treated (n=11) BRAF PTEN cohorts. Note that the BRAF PTEN vehicle- and AZD6244-treated cohort data are also used in Figs. 2 and 3. C) Left panel, representative H&E staining and IHC against phospho-ERK1/2 (Thr202/Tyr204), phospho-RPS6 (Ser235/Ser236), pRPS6 (Ser240/Ser244), cyclin D1, and p21 in BRAF PTEN mice treated with indicated treatment. Right panel, representative H&E staining and IHC against cyclin D1 and p21 in tumour sections from BRAF PTEN PREX2 mice treated with vehicle or AZD6244. D) Gene Set Enrichment Analysis (GSEA) plots showing downregulation of cell cycle-associated Hallmark gene sets, including E2F and MYC targets (V1), in xenografted BRAF PTEN tumours treated with AZD6244/AZD8186 combination *vs* AZD6244 monotherapy or AZD6244 monotherapy *vs* vehicle. E) Schematic of proposed pro-proliferative relationship between p110β and PREX2 in melanoma *in vivo*.

### Co-targeting of MEK1/2 and p110β has therapeutic efficacy in melanoma GEMMs

Having established the apparent relationship between PREX2 and p110β activity in PTEN-deficient melanoma both *in vitro* and *in vivo*, and demonstrated that targeting this relationship has significant translational potential. We went on to test whether it held true in mono- and combination therapy studies in immunocompetent, autochthonous melanoma GEMMs. As previously, BRAF PTEN or BRAF P53 mice, bearing a minimum of one cutaneous melanoma (diameter > 7 mm) (Supplementary Fig. S8E–F), were continuously treated to a defined endpoint (tumour size ≤ 15 mm, or ulceration). Despite resulting in a marginal, slowing of tumour growth over the first 7 days of treatment, AZD8186 monotherapy did not positively impact overall survival, nor did it prolong response in either the BRAF PTEN or BRAF P53 models (Fig. 6B Supplementary Fig. S8B). Compared with AZD6244 alone, AZD6244/AZD8186 had no additional impact on tumour regression in BRAF P53 mice, but it did provide a modest, yet significant, increase in overall survival and longevity of response relative to AZD6244 monotherapy (Fig. S7B). Moreover, combined treatment of BRAF PTEN mice with AZD6244/AZD8186 attenuated tumour growth, eliciting significant tumour regression over the first 7 days of treatment, and extended overall survival (Fig. 6B).

To understand whether the mechanisms associated with delayed onset of resistance in the BRAF PTEN model of melanoma correspond to those identified *in vitro*, we analysed the expression patterns of key biomarkers following 5 days of treatment in BRAF PTEN melanomas using immunohistochemistry. This indicated that whereas both AZD6244 and AZD6244/AZD8186 combination treatment equally suppressed MAPK signalling (pERK Thr202/Tyr204), RPS6 (Ser235/236) phosphorylation, and the expression and nuclear accumulation of cyclin D1 and p21, only RPS6 (Ser240/244) phosphorylation appeared more significantly suppressed by the combination treatment than AZD6244 monotherapy, at this timepoint (Fig. 6C). Moreover, the expression and nuclear accumulation of cyclin D1 and p21 were also suppressed in AZD6244-treated BRAF PTEN PREX2 melanomas (Fig. 6C – right panels).

Finally, the observed suppression of pro-proliferative signalling, inhibited growth both *in vitro* and *in vivo* is recapitulated by transcriptional profiling of BRAF PTEN tumours in response to short term treatment *in vivo*. As would be expected in a MAPK driven tumour model, MEK1/2 inhibition with AZD6244 resulted in dramatic remodelling of the transcriptional landscape, which was contrasted with the minimal impact of p110α (AZD8835) or p110β/δ (AZD8186) targeted monotherapy (Supplementary Fig. S8G). Critically, despite the lack of efficacy as monotherapy, both AZD8186 and AZD8835 amplified the transcriptional impact of AZD6244 (Supplementary Fig. 8G). Moreover, this specifically translates into an impact upon pro-proliferative gene signatures, with suppression of transcripts associated with activation of both E2F and cMyc transcriptional programmes following MEK1/2 inhibition, and deeper suppression of these same targets upon combined inhibition of MEK1/2 and p110β/δ (Fig. 6D).

In this study, we have used genetic deletion, or loss-of-GEF-function mutation, to model the effects of systemic inhibition of PREX2 signalling. We find that PREX2 loss or mutation strongly co-operates with MEK1/2 inhibition, in a manner phenocopied by inhibition or mutation of PIK3CB, to suppress growth of complex melanoma models, in a genotype-specific manner both *in vitro* and *in vivo*. Importantly, consistent with our findings, several studies have demonstrated that multiple PTEN-deficient tumour types are reliant on PI3Kβ/p110β activity, with PI3Kβ isoform-selective inhibition or genetic deletion of *PIK3CB –* but not *PIK3CA*/p110α – sufficient to perturb PI3K/AKT signalling and abrogate tumour cell growth *in vitro* and *in vivo*, with combination therapies showing the highest potency^37,38,43,44^. By contrast, recent studies have shown that melanomas harbouring *BRAF*V600E and *PTEN* deletion are refractory to PI3Kβ/p110β monotherapy blockade, with combined inhibition of PI3Kα/p110α, PI3Kγ/p110γ, and PI3Kδ/p110δ required to arrest their proliferation. Moreover, *RAC1*P29S-mutant melanoma cells have been recently shown to be refractory to p110β selective inhibition, albeit only *in vitro*, even though RAC1 activation directly regulates p110β^12^. Although the reasons for these discrepancies in drug vulnerabilities and dependencies remain unclear, it is important to note that our studies suggest that RAC1-activating mutations are mutually exclusive with *PTEN* loss, and that p110β inhibition selectively sensitizes cells to MEK inhibition in the subset of PTEN-deficient melanomas harbouring hyperactivation of RAC1 signalling, or broader dysregulation of the PREX2/RAC1/PI3Kβ axis. Nevertheless, together, these results suggest that combination therapy with isoform-specific inhibitors of PI3K, tailored to the patient’s profile, may offer a means to prolong the duration of the response to MAPK-targeted therapies in patients with *BRAF*-mutant, PTEN-deficient melanoma.

Despite almost unparalleled success in the use of targeted therapeutics for patient benefit in melanoma over the last two decades, and extensive research into the mechanisms underlying intrinsic and acquired therapy resistance, refractory or recurrent disease portends a dismal prognosis and remains a pressing problem in the clinic. Here, we demonstrate that genetic or therapeutic targeting of the PREX2/RAC1/p110β pathway can substantially enhance responses to MAPK targeting in *BRAF*-mutant melanoma both *in vitro* and *in vivo*, presumably through suppression of AKT/mTOR driven cell cycle progression (Fig. 6E). Given that clinically relevant p110β inhibitors are both currently available and well-tolerated, our research suggests a therapeutic approach which could be of significant benefit to patients in the future.

## Data Availability Statement

All data presented in this manuscript are archived on the Beatson server and are available on request.

## Competing interests

This work was funded in part through a research agreement with AstraZeneca. The funder had no role in the conceptualization, study design, data collection, analysis and interpretation, the decision to publish, or the preparation of the manuscript. O.J.S. receives funding from Boehringer Ingelheim, Novartis, and Cancer Research Horizons for research projects outwith the submitted work. S.T.B. is an employee of and shareholder in AstraZeneca PLC. The remaining authors declare no competing interests.

## Materials and Methods

### Mouse studies

All mouse studies were performed in accordance with UK Home Office regulations, under project licences 70/8646 and PP3908577, and were approved by the Animal Welfare and Ethical Review Board (AWERB) of the University of Glasgow. All mice were housed in conventional cages within a dedicated animal facility at a constant temperature (19–23 °C) and humidity (55 ± 10%), with a 12-h light/dark cycle and *ad libitum* access to food and water. The transgenic alleles used is this study were Tyr-Cre^ERT2 28^, *Braf*^LSL-V600E 45^, *Pten*^fl 46^, *Trp53*^fl 47^, *Prex2*^ko 48^, *Prex2*^E22A,N204A^ (designated *Prex2*^gd^) and *Pik3cb*^S205D,K224A^ (designated *Pik3cb*^rbd^) ^12^. All animal cohorts were maintained as C57Bl6 strains, typically inbred to a minimum of 4 generations. Genotyping for transgenic alleles and genetic background testing was carried out by Transnetyx (Cordova, TN, USA). For all mouse studies, no formal randomisation was performed but researchers were blinded to the mouse genotypes for analysis.

For melanoma GEMMs, genetic recombination was induced through daily topical administration of 2 mg tamoxifen (T5648, Sigma-Aldrich) on five consecutive days in mice of both sexes, aged 6 weeks–6 months. Mice were monitored by visual inspection for the development of melanocytic naevi and cutaneous melanoma at least twice weekly, with melanoma volume calculated as V=(length × width^2^)/2, where length is the greatest longitudinal diameter and width is the greatest transverse diameter, as measured by calipers. Tumour-bearing mice were sampled when exhibiting a cutaneous melanoma ≤15 mm in diameter, or tumour ulceration.

For xenograft studies, 1 × 10^6^ WM266.4 or A375 cells were resuspended in 100 µl of sterile phosphate-buffered saline (PBS) and subcutaneously engrafted into the flank of athymic CD1-*Foxn1^nu^* mice (Charles River). Tumour outgrowth was monitored by caliper measurements over time. Mice were aged to a humane endpoint defined by tumour size (diameter ≤15 mm) or ulceration.

### Generation of the Prex2***^gd^*** mouse strain

To generate a mouse strain with catalytically inactive (GEF-dead) PREX2, residues Glu^22^ and Asn^204^ in the catalytic DH domain were mutated to alanine using CRISPR/Cas9 gene editing. First, 20-nt sgRNAs were designed to direct wild type Cas9 to the relevant sites in exons 1 and 6 of *Prex2*, to be positioned directly upstream of a requisite 5’-NGG protospacer adjacent motif (PAM), and to have no highly homologous sites elsewhere in the mouse genome. Off-target potential was scored using software from the Feng Zhang laboratory (http://crispr.mit.edu/), Broad Institute (http://www.broadinstitute.org/rnai/public/analysis-tools/sgrna-design), and E-CRISPR (http://www.e-crisp.org/E-CRISP/). Three sgRNAs were selected for each target site. For exon 1, they were sgRNA 1 (GCGCTGAGCACGCACACGCGC-*AGG*), sgRNA 2 (GAAGACCGAGCGCGACTACG-*TGG*), and sgRNA 3 (GCGCGACTACGTGGGCACGC-*TGG*), with the adjacent PAM sequence in italics. For exon 6, they were sgRNA 4 (GCGTGTGTTCCAACATTAATG-*AGG*), sgRNA 5 (GTTGGAACACACGGCTTTCA-*TGG*) and sgRNA 6 (GTAATGAGGCCAAGAGACAGA-*TGG*). sgRNAs efficiency was assessed *in vitro* using the sgRNA *In vitro* Transcription and Screening kit (Clontech, 631439) following the manufacturer’s protocol, with 2 kb templates containing the target sequence amplified from mouse genomic DNA using primers GGTCAGTGGTGTGGTTGTTT and CCACCAAGTCCAGCTCAAAT for exon 1, and TCAGTTTTGAAATTGTGGTGCA and GCTGAGGGACATTCAAGACC for exon 6, respectively. All sgRNAs directed efficient cutting of the DNA by Cas9, which produced DNA fragments of the expected sizes. Two sgRNAs for each exon, namely sgRNAs 2 and 3 for exon 1, and sgRNAs 4 and 6 for exon 6, were selected for further assessment. Their efficacy in cells was tested using the Surveyor Mutation Detection Kit (Integrated DNA Technologies) following the manufacturer’s instructions. NIH/3T3 cells were transfected with pSpCas9(BB)-2A-GFP carrying the selected sgRNAs, genomic DNA was extracted, the relevant region amplified, annealed with wild type DNA, and treated with Surveyor nuclease to cut at the mismatches. Following that, sgRNA 3 was chosen for targeting exon 1 and sgRNA 6 for exon 6.

200 bp ssDNA repair templates were designed to introduce the desired point mutation by homology-directed repair, to introduce silent mutations creating restriction sites for screening purposes and, where possible, destroy the PAM sites, and to have symmetric homology arms of ≥90 nt. The exon 1 repair template was CTTGCCCCCAGCTCCGCGCCCCGCCGGCCACCATGAGCGACGAAAGCGCCAG GGAAGTAGACAAGCAGCTTCGCCTGCGCGTGTGCGTGCTCAGC**GCT**CTtCAGA AGACCGAGCGCGACTACGTGGGtACcCTaGAGTTCCTGGTGTCGGTGAGTAGCCGGCCCCGCGCACGGCACCAAGTCTGGAGCATTGTCTGC (nts coding for the E22A mutation in bold, silent mutations in lower case). The exon 6 repair template was TCTCCACAGGAATTACTGAAGCGGACTCCACGGAGACATAGTGACTACACAGCA GTGATGGAAGCACTCCAAGCCATGAAAGCCGTGTGTTCCAAtATT**GCT**GAGGCC AAGcGgCAaATGGAGAAACTGGAAGTTTTAGAAGAGTGGCAGGCACACATTGAAGGCTGGGAGGTACGTGTCCTTTGCTCAGCTCTT (nts coding for the N204A mutation in bold, silent mutations in lower case). These ssDNAs were purchased from Dharmacon as PAGE-purified Ultramer ssDNAs.

The selected sgRNA, ssDNA repair template and Cas9 mRNA were microinjected into the pronucleus of C57BL/6J mouse zygotes by the Babraham Gene Targeting Facility. Initially, mice carrying the E22A mutation were generated. Pups were genotyped by sequencing of 571 bp PCR products amplified from genomic DNA using GACTGTCCCGTTCTGAGTCC forward and AATTTGCCCTGGGAGATGGA reverse primers. Heterozygous Prex2^E22A/+^ mice were bred together to generate homozygous Prex2^E22A/E22A^ animals, which were then subjected to a further round of pronuclear injections to target the second site, Prex2^N204A^. Pups were genotyped for the N204A mutation by sequencing of 536 bp PCR products, amplified from genomic DNA using TGCTCACTCATGGATTTGACC forward and TCCATCACACATGTCTCAGGT reverse primers. Prex2^E22A/E22A;N204A/+^ mice were bred together to generate homozygous double knock-in Prex2^gd^ (Prex2^E22A/E22A;N204A/N204A^) mice, which were born at the expected Mendelian rate, were fertile, bred well and appeared healthy. Once the Prex2^gd^ strain was fully established, routine genotyping was done by Transnetyx (Cordova, TN, United States).

### In vivo therapeutic studies

Clinical-grade AZD6244 ^49^, AZD8186 ^38^, AZD8835 , AZD2014 , and AZD5363 were supplied by AstraZeneca under a collaborative research agreement. For *in vivo* studies, AZD6244 and AZD8186 were prepared in a vehicle of 0.5% (w/v) hydroxypropyl methylcellulose (#09963, Sigma-Aldrich) and 0.1% (v/v) Tween-80 (#P8192, Sigma-Aldrich) in water, and administered at a dose of 25 mgkg^-1^ and 50 mgkg^-1^, respectively, in 100 μl. For combination-dosing studies, compounds were co-formulated in the same vehicle and administered with the same dose and volume as monotherapy. For cutaneous melanoma and xenografted tumour treatment studies, mice were enrolled onto vehicle, mono- or combination treatment in a pseudo-randomised manner when exhibiting a cutaneous melanoma >7 mm in diameter. Mice were excluded from study where tumour ulceration was observed within 7 days of enrolment. Therapeutic response was measured both in terms of primary tumour size (growth/regression) and overall survival.

### Transcriptional profiling by RNAseq

Melanoma fragments isolated from BRAF PTEN donor mice were maintained as xenografts via subcutaneous implantation into the flank of adult athymic CD1-*Foxn1^nu^*. P1 tumour fragments were then implanted subcutaneously into the flank of C57Bl6/J recipients, with these recipient mice were enrolled onto treatment when exhibiting a single tumour with a diameter >7mm. Therapeutic treatments were as outlined in the in vivo therapeutic studies section above, but sacrificed following 3 days of treatment. RNA was isolated using the Qiagen RNAeasy mini kit (#74104), in accordance with the manufacturer’s instructions. Tumour tissue was lysed using a Precellys Lysing Kit (#KT03961-1-003-2) and Precellys Evolution tissue homogeniser from Bertin Technologies. RNA quality was assessed using an Agilent 2200 Tapestation, with RNA screen tape. Libraries for cluster generation and DNA sequencing were prepared using the Illumina TruSeq RNA LT Kit. DNA library quantity and quality was assessed on a Agilent 2200 Tapestation (D1000 screentape) and Qubit (Thermo Fisher Scientific) respectively. Libraries were sequenced using the Illumina Next Seq 500. Gene set enrichment analysis (GSEA) was employed using fgsea R package (v1.21.0), using a ranked gene list via limma R package (v3.50.3) in a grouped pairwise manner. Statistical significance was measured with Benjamini-Hochberg (BH) False Discovery Rate (FDR) < 0.05 and normalised enrichment score (NES) indicates the upregulation (positive value) and down-regulation (negative value).

### Histochemical and immunohistochemical staining

Haematoxylin and eosin (H&E) staining and immunohistochemistry (IHC) were performed on 4-µm formalin-fixed paraffin-embedded (FFPE) sections which had previously been baked at 60 °C for 2 h.

Staining was performed in a Leica Biosystems BOND RX autostainer, using antibodies against: cyclin D1 (#55506, Cell Signaling), Ki67 (#12202, Cell Signaling), phospho-RPS6 Ser240/Ser244 (#5364, Cell Signaling), and p21 (#ab107099, Abcam). FFPE sections were deparaffinized using BOND Dewax solution (#AR9222, Leica Biosystems) and epitope retrieval using ER2 solution (#AR9640, Leica Biosystems) for 20 min at 95 °C, except for cyclin D1, where sections underwent retrieval for 30 min. Sections were rinsed with BOND Wash Solution (#AR9590, Leica Biosystems) before endogenous peroxidase blocking using a BOND Intense R Detection kit (#DS9263, Leica Biomarkers) for 5 min. Typically, for IHC, sections were rinsed with Wash Solution before application of a blocking solution from an anti-rat ImmPRESS Detection kit (#MP7444-15, Vector Labs) for 20 min. Sections were then rinsed with Wash Solution before application of primary antibodies at an optimised dilution (cyclin D1, 1:150; Ki67, 1:1000; RPS6 Ser240/Ser244, 1:1000; p21, 1:250) for 30 min. Sections were then rinsed with Wash Solution and incubated for 30 min with anti-rabbit EnVision+ HRP-labelled polymer secondary antibody (K4003, Agilent), except for p21 where an anti-rat ImmPRESS secondary antibody was applied. Sections were rinsed with wash buffer and visualised using 3,3′-diaminobenzidine (DAB) from the BOND Intense R Detection kit.

IHC staining for phospho-RPS6 Ser235/Ser236 (#4858, Cell Signaling) and phospho-p44/42 MAPK (ERK1/2; ERK1, Thr202/Tyr204/ERK2,Thr185/Tyr187) (#9101, Cell Signaling) was performed on an Agilent Autostainer Link 48. Sections were loaded into an Agilent pre-treatment module, deparaffinised and subjected to heat-induced epitope retrieval (HIER), at 97 °C for 20 min, using EnVision FLEX target retrieval solution, high pH (#K8004, Agilent). After HIER, sections were washed thoroughly with EnVision FLEX Wash Buffer (#K8007, Agilent), loaded onto the autostainer, subjected to endogenous peroxidase blocking (#S2023, Agilent) for 5 min, and rinsed with FLEX Wash Buffer. Primary antibodies were applied at an optimized dilution (phospho-RPS6 Ser235/Ser236, 1:75; phospho-p44/42 MAPK, ERK1/2, 1:400) for 30 min at room temperature. Sections were then rinsed with FLEX Wash Buffer, incubated with anti-rabbit EnVision secondary antibody for 30 min, and rinsed with FLEX Wash Buffer. Staining was visualised with Liquid DAB+ (#K3468, Agilent). After staining for 10 min, sections were washed in tap water and counterstained with haematoxylin “Z” stain (#RBA-4201-00A, CellPath).

H&E staining was performed with a Leica autostainer (#ST5020). Sections were dewaxed, rehydrated through graded alcohols, stained with haematoxylin “Z” stain (#RBA-4201-00A, CellPath) for 13 min, washed in tap water, differentiated in 1% (v/v) acid alcohol, washed, with nuclei blued in Scott’s tap water substitute. After further washing, the sections were stained with Putt’s Eosin for 3 min.

After H&E staining or IHC, sections were rinsed in tap water, dehydrated through graded ethanols, and placed in xylene. The stained sections were coverslipped in xylene using DPX mountant (#SEA-1300-00A, CellPath).

### Cell culture

WM266.4, A375, WM793, WM1158, and A2058 cells were the kind gift of Prof. Lionel Larue (Institut Curie, Orsay, France). WM266.4, A375, WM1158, and A2058 were maintained in DMEM (#21969, Thermo Fisher Scientific) supplemented with 10% (v/v) foetal calf serum (FCS) (#10270, Thermo Fisher Scientific), 200 μM L-glutamine (#25030, Thermo Fisher Scientific) and 100 U/ml^-1^ penicillin/streptomycin (#15140, Thermo Fisher Scientific). WM793 were maintained in RPMI 1640 (#31870, Thermo Fisher Scientific) medium supplemented with 10% (v/v) FCS, 200 µM L-glutamine and 100 U/ml^-1^ penicillin/streptomycin (#15140, Thermo Fisher Scientific). Cells were harvested by trypsinization with 0.25% (w/v) trypsin (#15090, Thermo Fisher Scientific), washed with PBS, and centrifuged. Cell pellets were resuspended in the appropriate culture medium. The enzymatic activity of trypsin was blocked by resuspension in complete medium, and cell suspensions were centrifuged, washed in PBS, resuspended in the appropriate culture medium.

For *in vitro* drug treatments, all inhibitors were prepared in dimethyl sulphoxide (DMSO; Sigma-Aldrich) as a 1000× stock and diluted into the appropriate buffer/medium to the indicated final concentration – AZD6244, 100 nM; AZD8186, 250 nM; AZD8835, 250 nM; AZD2014, 500 nM; AZD5363, 250 nM.

### Cell proliferation

WM266.4 (2 × 10^3^), A375 (1.3 × 10^3^) or WM793 (1.5 × 10^3^) cells, were seeded into 96-well plates for 24 h before addition of AZD6244 (100 nM), AZD8186 (250 nM), AZD8835 (250 nM), AZD2014 (500 nM), AZD5363 (250 nM), a combination of these drugs or DMSO for up to 96 h. Growth of cultures was monitored at 6-h intervals for up to 96 h using an IncuCyte live-cell analysis system (Essen Bioscience). IncuCyte software was used to measure relative cell confluence, which was normalised to starting confluence. Experiments were performed independently at least 3 times with technical triplicates.

### Immunoblotting

WM266.4, A375 or WM793 cells were seeded (2 x 10^5^ cells) in media (DMEM for WM266.4 and A375 and RPMI for WM793) supplemented with 10% FBS, 2mM glutamine + 100Uml^-1^ penicillin/streptomycin, into 6-well plates for 24 hr before culturing in AZD6244 (100 nM), AZD8186 (250 nM), AZD8835 (250 nM), AZD2014 (500 nM), AZD5363 (250 nM) or a combination of these drugs for 24 hr. Cells were harvested & lysed using RIPA with protease inhibitors (Roche 11836153001) & phosphatase inhibitors (Roche 04906845001). 20 μg protein was loaded onto a 4-12% bis-tris gradient protein gel (Invitrogen) under reducing conditions and transferred onto PVDF membrane. Membranes were blocked in 5% milk/TBST, probed overnight with primary antibodies diluted in 5% BSA/TBST followed by suitable HRP-conjugated secondary antibodies diluted in 5% milk/TBST. Primary antibodies from Cell Signaling Technology were phospho-p90RSK (Ser380) 9341, phospho-Akt (Ser473) 3787, PTEN 9559, phospho-Erk1/2 (Thr202/Tyr204) 9101, phospho-S6 ribosomal protein (Ser235/Ser236) 2211, phospho-S6 ribosomal protein (Ser240/Ser244) 5364, phospho-4E-BP1 (Thr37/Thr40) 2855, phospho-Rb (Ser793) 9301 and cyclin D1 2978 and beta-actin antibody was from Sigma A2228. Blots were developed using Clarity Western ECL and ChemiDoc Imaging System (Bio-Rad).

### Reverse-phase protein array

WM266.4, WM793, WM1158, A375 and A2058 cells treated with AZD6244, AZD8186, AZD8835 or combinations of these were washed with PBS, then lysed in a buffer comprised of 1% Triton X-100, 50 mM HEPES (pH 7.4), 150 mM sodium chloride, 1.5 mM magnesium chloride, 1 mM EGTA, 100 mM sodium fluoride, 10 mM sodium pyrophosphate, 1 mM sodium vanadate and 10% (v/v) glycerol, supplemented with cOmplete ULTRA and PhosSTOP protease and phosphatase inhibitor cocktails (Roche). Following clearing by centrifugation, lysates were diluted to produce a dilution series of each sample, and spotted onto nitrocellulose-coated slides (Grace Bio-Labs) in triplicate using an Aushon 2470 array platform (Aushon Biosystems). Slides were then blocked in SuperBlock (TBS) blocking buffer (Thermo Fisher Scientific) and incubated with validated primary antibodies (1:250, Supplementary Table S2). Bound antibodies were detected by incubation with DyLight 800-conjugated secondary antibodies (New England BioLabs), and analysed using an InnoScan 710-IR scanner (Innopsys). The relative fluorescence intensity of each array feature was quantified using Mapix software (Innopsys). Intensity values were normalised to the DMSO control samples for each cell line, and log_2_ transformed data subsequently plotted.

### *In vitro* cell-cycle analysis

WM266.4 or A375 cells were seeded at 4 × 10^5^ cells/6-cm petri dish in DMEM supplemented with 10% (v/v) FBS, 2 mM glutamine, and 100 U/ml^-1^ penicillin/streptomycin. After 24 h, cells were synchronised by culturing in serum-free medium for a further 24 h. Synchronised cultures were then treated for 24 h with AZD6244 (100 nM), AZD8186 (250 nM), a combination of AZD6244 (100 nM) and AZD8186 (250 nM) (AstraZeneca), or DMSO, in DMEM containing 10% FBS, 2 mM glutamine, and 100 U/ml^-1^ penicillin/streptomycin. Culture supernatants and trypsinised cells were harvested, and pelleted cells were fixed in ice-cold 70% ethanol. Fixed cells were stained with FxCycle PI/RNase staining solution (#F10797, Thermo Fisher Scientific) as per manufacturer’s instructions and acquired on the Attune Flow Cytometer (Thermo Fisher Scientific) followed by analysis using FlowJo software (BD). Experiments were performed independently at least 3 times.

### CRISPR/Cas9-mediated genome editing

Genome editing was performed in WM266.4 cells. Alt-R™ S.p. Cas9 Nuclease V3, 100 µg (#1081058), Cas9 Electroporation Enhancer (#1075915), tracrRNA, and guides were purchased from Integrated DNA Technologies (IDT) and prepared according to the manufacturer’s instructions. Electroporation was performed with the Lonza SF Cell line 4D-Nucleofector X solution (#V4XC-2032) and the 4D-Nucleofector X Unit (Lonza) (#AAF-1003X) using program CM137. To validate the CRISPR knockout, gene-edited sequences were PCR-amplified and sequenced (in-house Molecular Technology Service) and knockout efficiency was quantified using the decomposition algorithm TIDE^42^. Targeting gRNAs were designed by, and purchased from Integrated DNA Technologies (IDT) as follows:

*PREX1* 5’-GCTATACCGTCACCAACGGCTGG-3’

*PREX2* 5’-TCGTGGCCGGATCAACACGGAGG-3’

*PIK3CB* 5’-CTTCCCGAGGTACCTCCAACTGG-3’

Primers used for coding region amplification were as follows:

*PIK3CB* Forward: 5’-TCCTTGACATCTGGGCGGTGGA-3’

*PIK3CB* Reverse: 5’-AGGCAAGCCTGCTGAGGGAAAA-3’

*PREX1* Forward: 5’-GCCCAGGAAGCATTTTGGGGCT-3’

*PREX1* Reverse: 5’-TGCCCCTTCCCTAGACACAGCC-3’

*PREX2* Forward: 5’-CAGAGTCTGATTGGGCACCGCT-3’

*PREX2* Reverse: 5’-TCACAGTAGTCCTCCCCTCCCT-3’.

### Statistical analysis and data visualisation

Statistical analysis and graph plotting was carried out using GraphPad Prism (10.0.2) (LaJolla, USA). All comparisons made, and statistical tests used are described in the appropriate figure legend.

## Acknowledgements

The research outlined in this manuscript was made possible through a collaborative research agreement with AstraZeneca and funding to C.A.F., A.D.C. and O.J.S. In addition, C.A.F., O.J.S. and A.D.C. received funding from the Cancer Research UK Grand Challenge Rosetta Consortium (A25045), D.K. and M.F. were funded by EU Horizon 2020 Sys-mel Consortium (611107), P.P.C. and K.G. were funded by the Cancer Research UK Grand Challenge SpecifiCancer Consortium (A29055), S.K.F. was funded by a Pancreatic Cancer UK Future Leaders Fellowship awarded to O.J.S. and J.P.M., and C.F., D.K., P.P.C., M.F., S.K.F., N.V., K.G., C.N., J.P.M, O.J.S and A.D.C. were supported by CRUK Beatson Institute core funding (A17196, A31287). The authors would like to acknowledge Dr. Joanna Birch (University of Glasgow) for critical review of the manuscript.

## Author Contributions

C.A.F., D.K., P.P.C., M.F., O.J.S. and A.D.C., designed the research and interpreted the data. C.A.F., D.K., M.F., E.T., N.V., V.P., and A.D.C. performed experiments and analysed data. B.S., A.F.M., and J.C.D. performed and analysed RPPA analysis. A.D.C., K.G. and C.B. performed transcriptomics and analysed transcriptomic data. C.N., N.O.C., D.C.H., P.D.D, J.D., H.C.E.W. and S.T.B. provided reagents and advice. N.S., O.J.S., and A.D.C. wrote the paper. All authors contributed to the manuscript.

## Supplementary Figures

**Supplementary Fig. S1:**
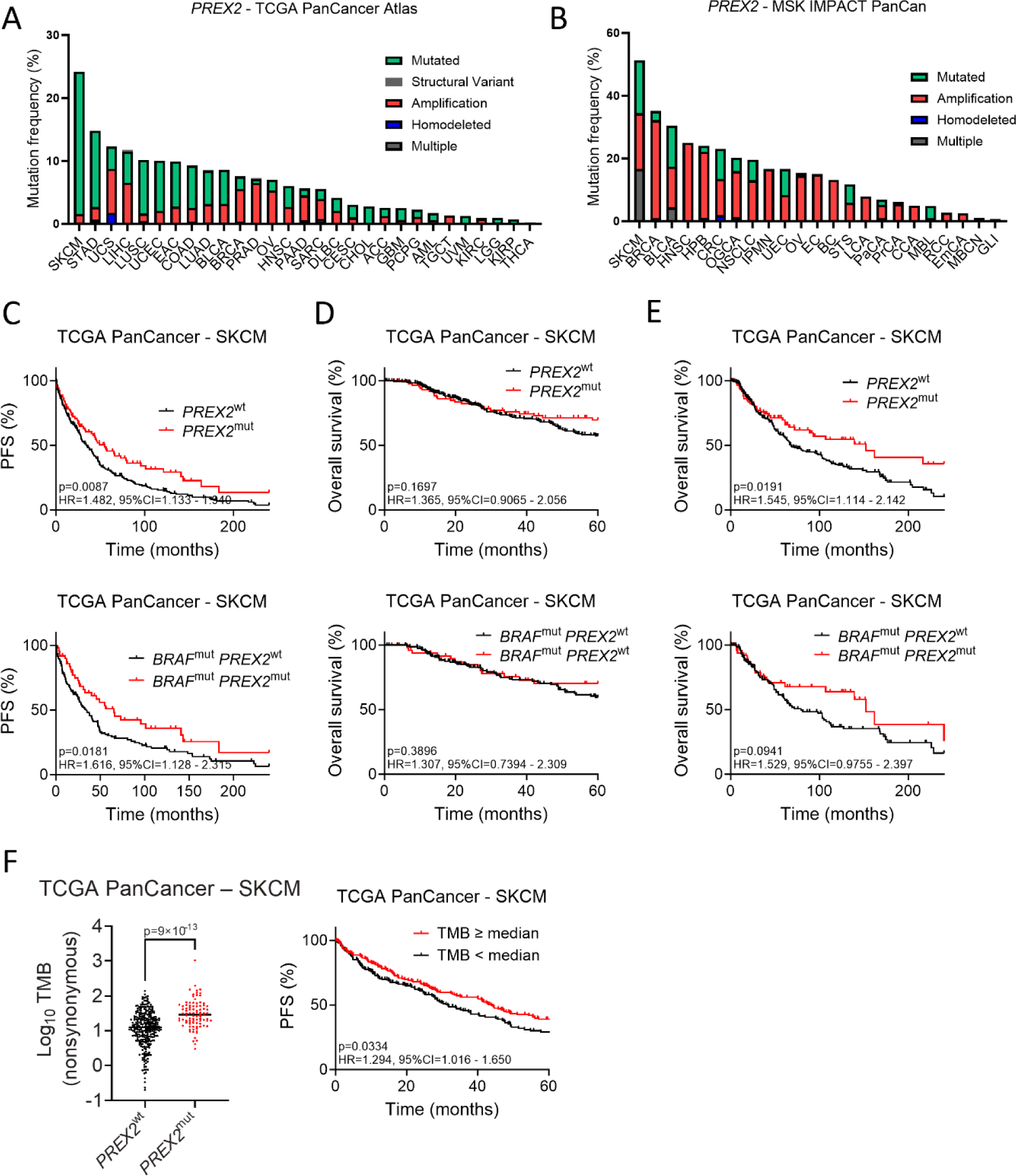
Impact of *PREX2* mutation in human cancers. A) *PREX2* mutation frequency and type across indicated cancers in/from the TCGA PanCancer Atlas patient cohort. B) *PREX2* mutation frequency and type across indicated cancer types in the MSK-IMPACT PanCancer patient cohort. Please see Supplementary Table S1 for abbreviations. C) Progression-free survival (PFS) of *PREX2* mutant *vs* wild-type cases in the curated cohort of SKCM patients from the TCGA PanCancer cohort, censored at 20 years. Top panel, all cases (n=381; *PREX2* wild-type = 290, *PREX2* mutant = 91); Bottom panel, *BRAF*-mutant cases (n=207; *PREX2* wild-type = 155, *PREX2* mutant = 52). D) Overall survival of *PREX2*-mutant *vs* wild-type SKCM cases from the TCGA PanCancer Atlas curated patient cohort, censored at 5 years. Top panel, all cases (n = 380; *PREX2* wild-type = 289, *PREX2* mutant = 91); Bottom panel, *BRAF*-mutant cases (n = 206; *PREX2* wild-type = 154, *PREX2* mutant = 52). E) Overall survival of *PREX2*-mutant *vs* wild-type SKCM cases from the TCGA PanCancer Atlas curated patient cohort, censored at 20 years. Top panel, all cases (n = 380; *PREX2* wild-type = 289, *PREX2* mutant = 91); Bottom panel, *BRAF*-mutant cases (n = 206; *PREX2* wild-type = 154, *PREX2* mutant = 52). F) Left panel, total mutation burden (TMB) of *PREX2*-mutant (n = 91) *vs* wild-type (n = 290) SKCM cases from the TCGA PanCancer Atlas curated patient cohort. TMB was calculated by the number of non-synonymous mutations, comprising single nucleotide variants, splice-site variants, and short insertions and deletions (InDels), per Mb of coding regions. Centre line, median TMB. Right panel, progression-free survival (PFS) of SKCM patients stratified by median TMB, censored at 5 years. TMB<median, n = 190, TMB≥median, n = 191.

**Supplementary Fig. S2:**
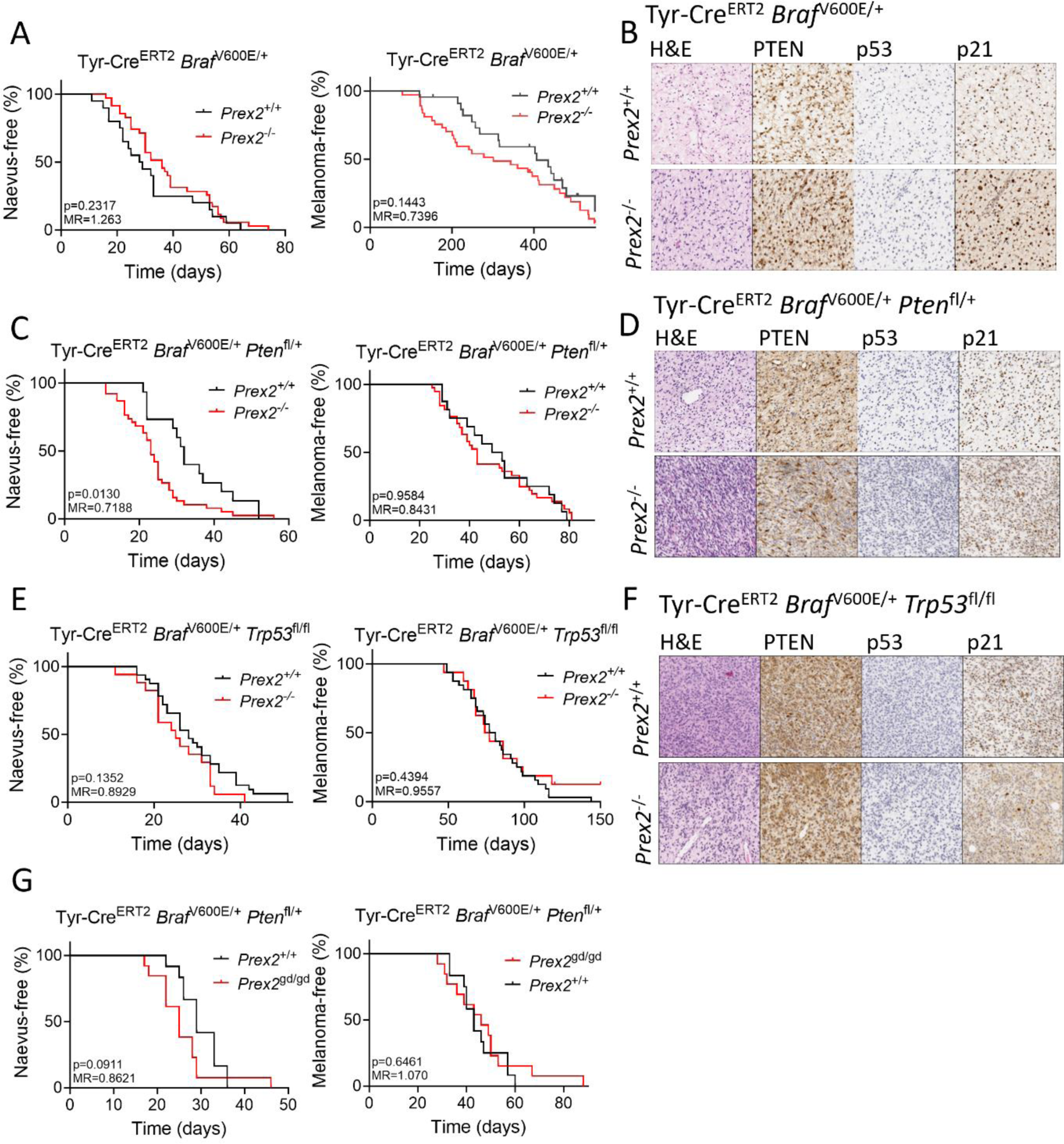
Characterisation of the disease trajectory and phenotype of PREX2-deficient melanoma models. A) Naevus-free (left) and melanoma-free (right) survival of BRAF and BRAF PREX2 cohorts. Tick marks indicate melanoma-free mice censored at indicated times post induction. B) H&E staining and IHC against PTEN, p53, and p21 in tumours from mice in (A). C) Naevus-free (left) and melanoma-free (right) survival of BRAF PTEN and BRAF PTEN PREX2 cohorts. D) H&E staining and IHC against PTEN, p53, and p21 in tumours from mice in (C). E) Naevus-free (left) and melanoma-free (right) survival of the indicated BRAF P53 and BRAF P53 PREX2 cohorts. F) H&E staining and IHC against PTEN, p53, and p21 in tumours from mice in (E). G) Naevus-free (left) and melanoma-free (right) survival of BRAF PTEN and BRAF PTEN PREX2-GD cohorts. MR, median ratio.

**Supplementary Fig. S3:**
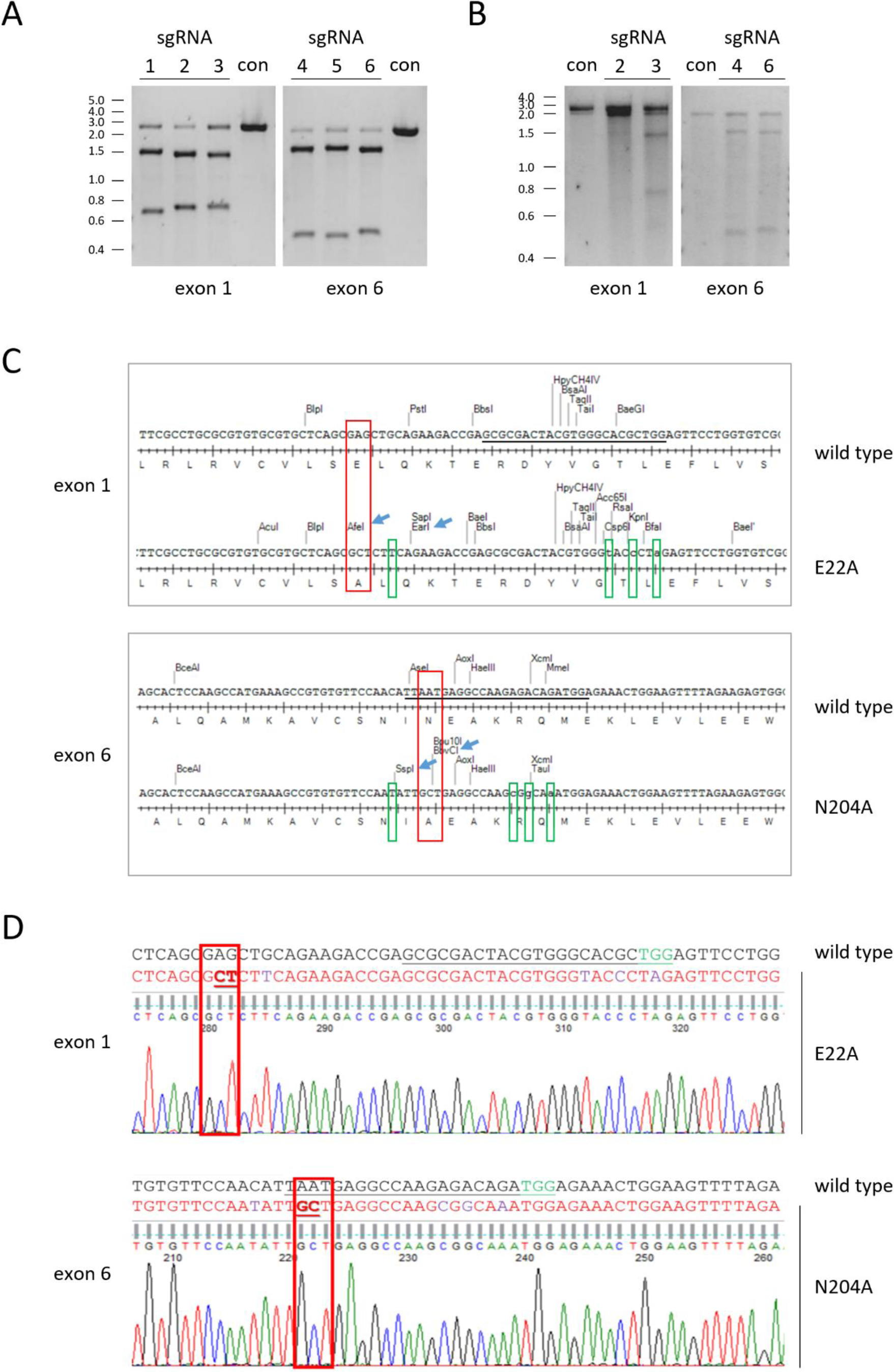
Targeting strategy for catalytically inactive *Prex2* transgenic allele. (A) Assessment sgRNA efficiencies *in vitro*. PCR products of *Prex2* mouse genomic sequence encompassing the target sites in exon 1 and exon 6 were cleaved by recombinant Cas9 nuclease in the presence of candidate sgRNAs, as indicated. Control DNA fragments were mock-treated in absence of sgRNA. (B) Assessment sgRNA efficiency in cells. NIH/3T3 cells were transfected with sgRNAs and Cas9 to target sites in exon 1 and exon 6 of *Prex2*. The relevant regions of genomic DNA were amplified, annealed with wild type DNA, and treated with Surveyor nuclease to cut at the mismatches. Wild type DNA fragments were used as controls. (C) Restriction maps of part of the DNA repair templates used to introduce point mutations in *Prex2* exons 1 and 6, compared to the wild type sequence. Red boxes show the nucleotide changes which result in the E22A and N204A mutations. Green boxes highlight silent mutations introduced to create restriction sites or destroy PAM motifs. Blue arrows show restriction enzyme sites useful for screening. (D) Representative sequencing traces of a homozygous *Prex2*^gd^ (*Prex2*^E22A/ E22A;N204A/N204A^) mouse (red letters) compared to wild type (black letters). The sgRNA sequences are underlined in black, the PAM sites in green. Red boxes show the nucleotide changes introduced to render the protein catalytically inactive. Purple letters show silent point mutations introduced to create restriction enzyme sites and/or destroy PAM motifs.

**Supplementary Fig. S4:**
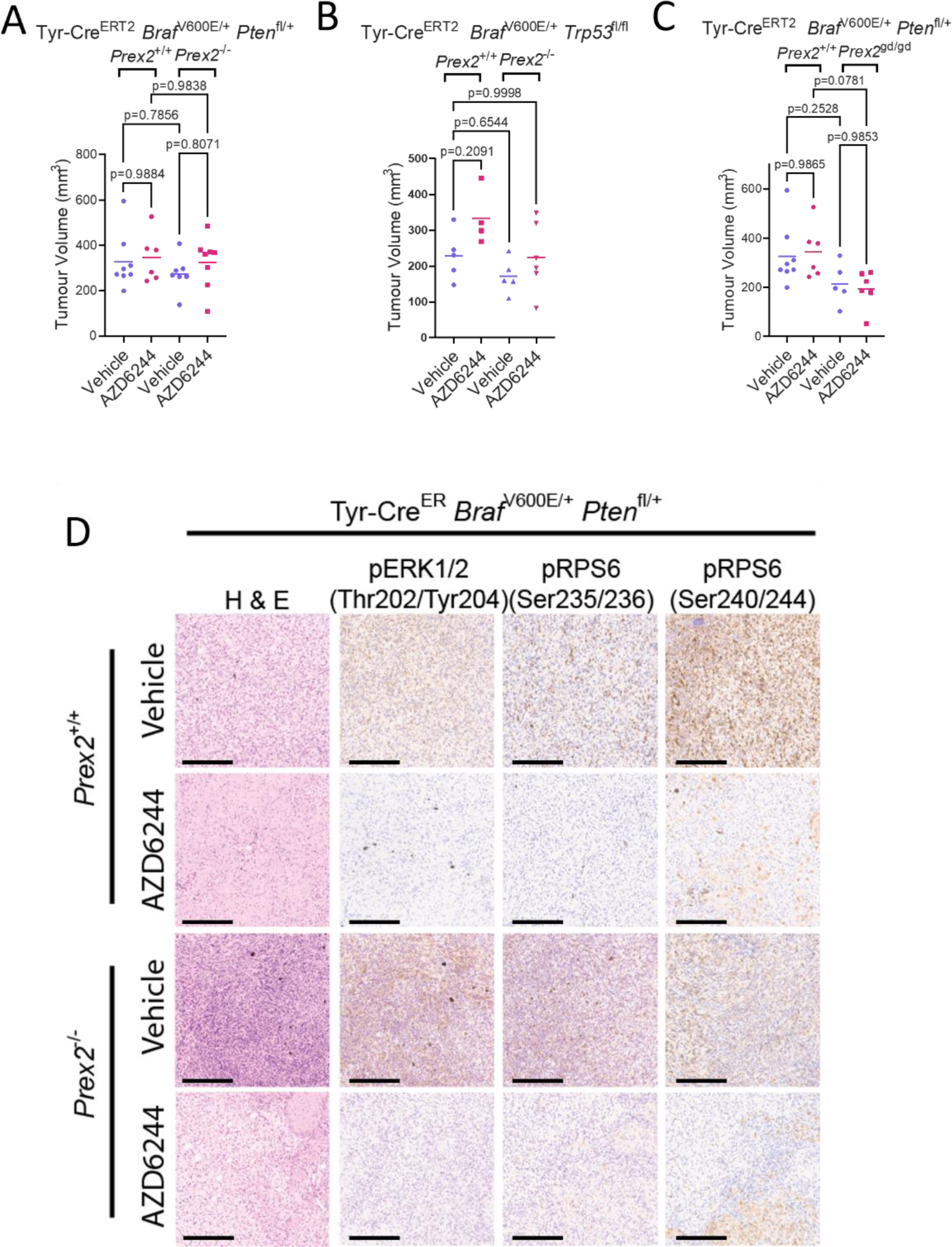
Therapeutic impact of MAPK inhibition in PREX2-deficient melanoma. A–C) Pre-treatment tumour volume in BRAF PTEN (vehicle, n=8; AZD6244, n=6) *vs* BRAF PTEN PREX2 (vehicle, n=7; AZD6244, n=8) (B), BRAF P53 (vehicle, n=5; AZD6244, n=4) *vs* BRAF P53 PREX2 (vehicle, n=5; AZD6244, n=6) (C), and BRAF PTEN (vehicle, n=8; AZD6244, n=6) *vs* BRAF PTEN PREX2-GD (vehicle, n=5; AZD6244, n=6) (D) cohorts. Note that the same BRAF PTEN treatment datasets are represented in B and D. p-values calculated by one-way ANOVA corrected for multiple comparisons (Tukey). D) Representative H&E staining and IHC against phospho-ERK1/2 (Thr202/Tyr204), phospho-RPS6 (Ser235/236), and phospho-RPS6 (Ser240/244) in tumours from BRAF PTEN and BRAF PTEN PREX2 cohorts following 5-day treatment with AZD6244 or vehicle. Scale bar – 200 µm.

**Supplementary Fig. S5:**
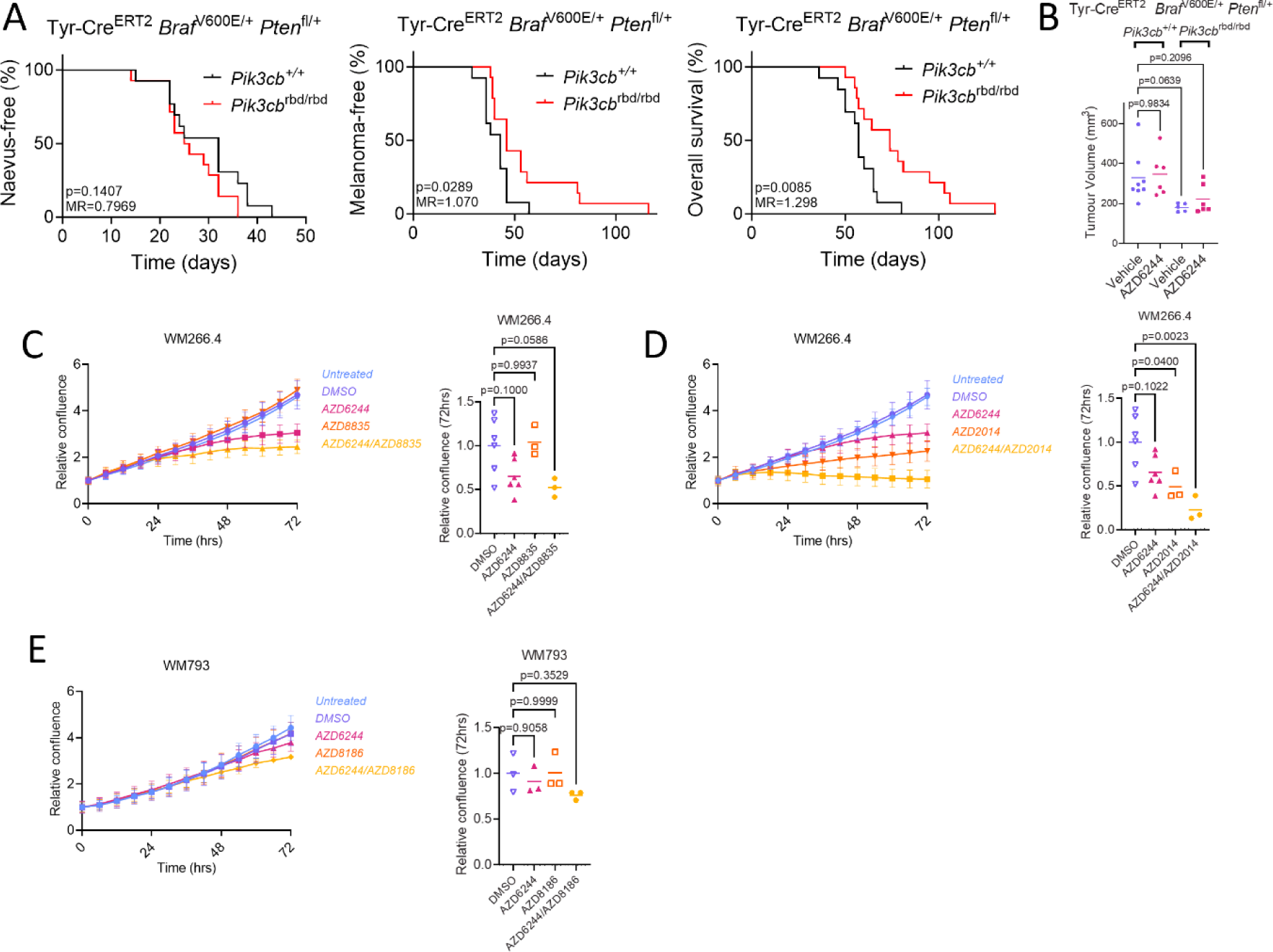
Targeting p110β function in melanoma as a therapeutic opportunity. A) Naevus-free (left), melanoma-free (centre), and overall (right) survival of BRAF PTEN *vs* BRAF PTEN PIK3CB cohorts. MR, median ratio. p-values calculated by log-rank (Mantel–Cox) test. B) Pre-treatment tumour volume in BRAF PTEN *vs* BRAF PTEN PIK3CB cohorts. BRAF PTEN+vehicle, n=8; BRAF PTEN+AZD6244, n=6; BRAF PTEN PIK3CB*+*vehicle, n=4; BRAF PTEN PIK3CB*+*AZD6244, n=6. p-values calculated by one-way ANOVA corrected for multiple comparisons (Tukey). C) Left panel, *in vitro* longitudinal growth of WM266.4 cells treated with mono- or combination therapy comprising AZD6244 (MEK1/2 inhibitor) and/or AZD8835 (p110α inhibitor). Right panel, confluence of WM266.4 cells, treated with indicated treatments, relative to starting confluence at 72 h. Vehicle, n=6; AZD6244, n=6; AZD8835, n=3; AZD6244/AZD8835, n=3. p-values calculated by one-way ANOVA corrected for multiple comparisons (Tukey). D) Left panel, *in vitro* longitudinal growth of WM266.4 cells treated with mono- or combination therapy with AZD6244 (MEK1/2 inhibitor) and/or AZD2014 (mTOR inhibitor). Right panel, confluence of WM266.4 cells, treated with indicated treatments, relative to untreated control at 72 h. Vehicle, n=6; AZD6244, n=6; AZD2014, n=3; AZD6244/AZD2014, n=3. p-values calculated by one-way ANOVA corrected for multiple comparisons (Tukey). Note that the same vehicle and AZD6244 treated datasets are presented in C and D. E) Left panel, *in vitro* longitudinal growth of WM793 cells treated with mono- or combination therapy with AZD6244 (MEK1/2 inhibitor) and/or AZD8186 (p110β/δ inhibitor, 250 nM). Right panel, confluence of WM793 cells, treated with indicated treatments, relative to untreated control at 72 h. Vehicle, n=3; AZD6244, n=3; AZD8186, n=3; AZD6244/AZD8186, n=3. p-values calculated by one-way ANOVA corrected for multiple comparisons (Tukey). Note that the same vehicle and AZD6244 treated datasets are presented in both panels. In (C–E) left panels, data represent mean ± SEM (relative cell confluence at each timepoint). In (B) and (C–E) right panels, the centre line represents the mean.

**Supplementary Fig. S6:**
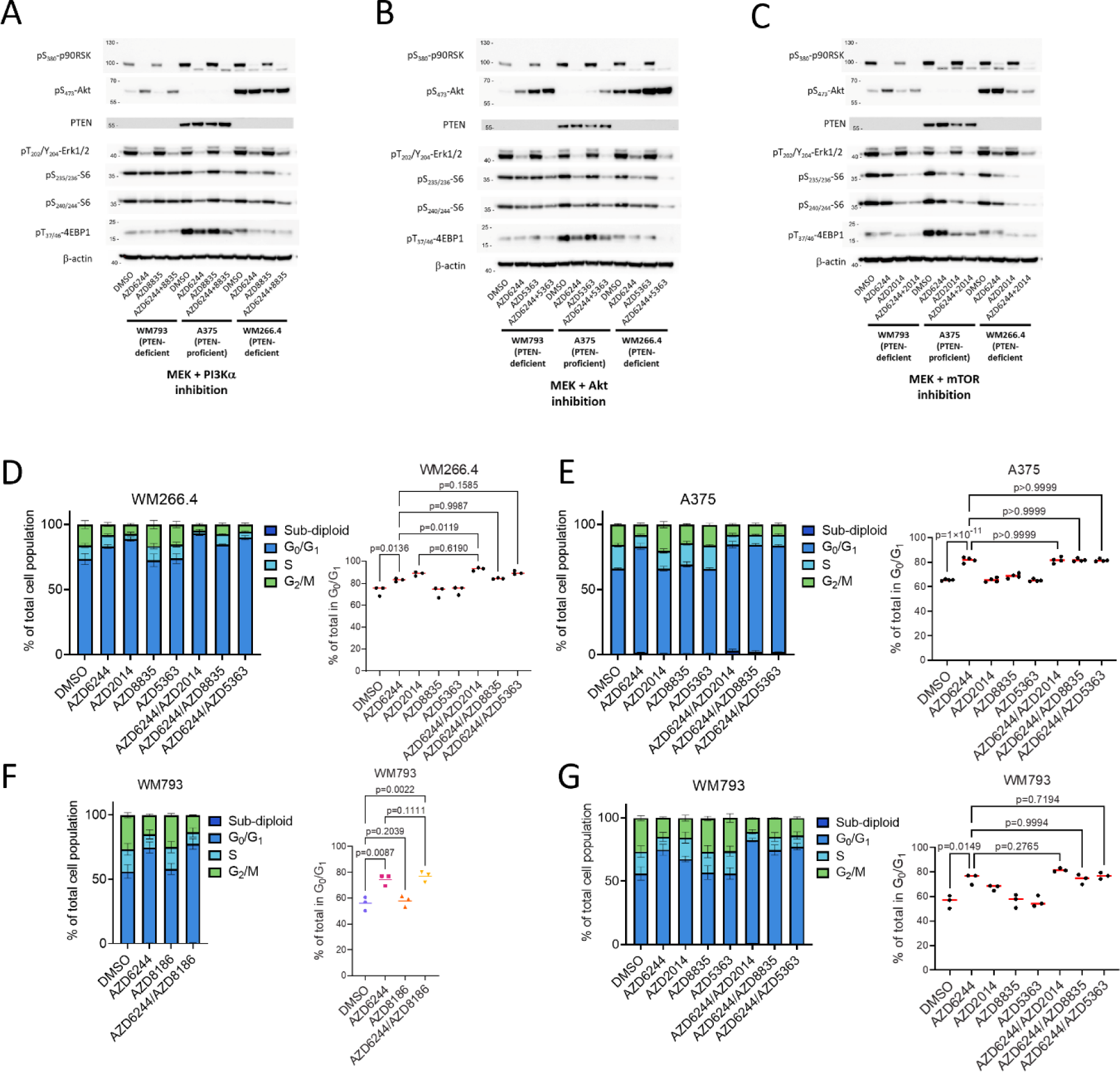
Inhibitors of AKT or mTOR, but not p110α, cooperate with MEK inhibition and phenocopy p110β/MEK co-targeting. A–C) Immunoblotting for indicated activated components of the MAPK–PI3K–mTOR pathways in WM266.4, A375, and WM793 human melanoma cells treated with indicated treatments. β-actin serves as a sample integrity control. The blots are representative of 3 repeated experiments. D) Left panel, flow cytometry–based cell-cycle profiling of WM266.4 cells following indicated 24 h treatment. Right panel, proportion of treated WM266.4 cells in G_1_/S at 24 h. n=3 per treatment. Statistical testing by one-way ANOVA corrected for multiple comparisons (Tukey). E) Left panel, flow cytometry–based cell-cycle profiling of A375 cells following indicated 24 h treatment. Right panel, proportion of treated A375 cells in G_1_/S at 24 h. n=4 per treatment. Statistical testing by one-way ANOVA corrected for multiple comparisons (Tukey). F,G) Left panels, Flow cytometry–based cell-cycle profiling of WM793 melanoma cells following indicated 24 h treatment. n=3 per treatment. Right panels, proportion of treated WM793 cells in G_1_/S at 24 h. Statistical testing by one-way ANOVA corrected for multiple comparisons (Tukey).

**Supplementary Fig. S7:**
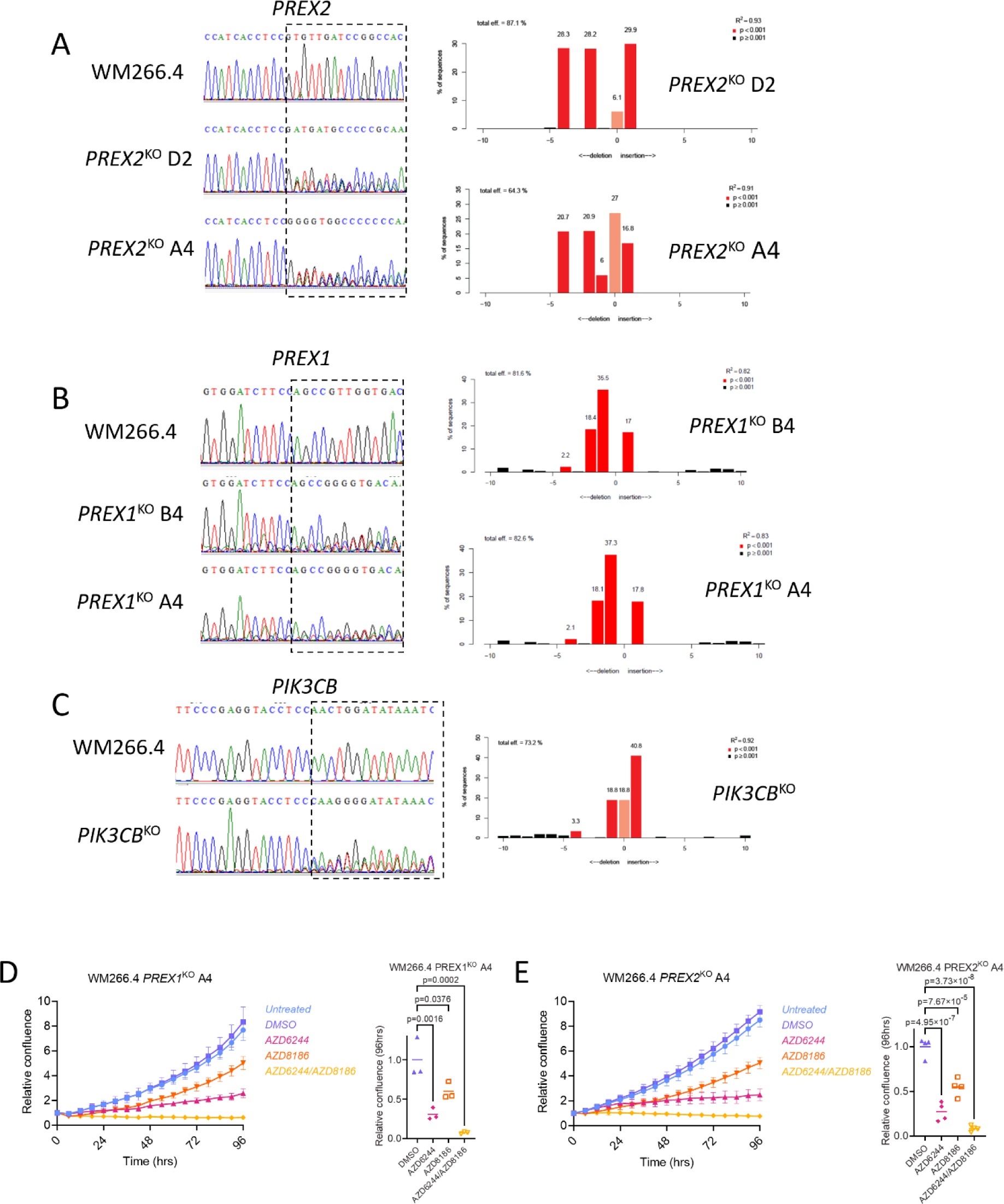
CRISPR/Cas9-mediated targeted deletion of *PREX1, PREX2,* and *PIK3CB* in WM266.4 cells *in vitro*. A) Left panel, Sanger sequencing trace depicting region immediately upstream and downstream of the targeted protospacer-adjacent motif (PAM) site in *PREX2* in parental WM266.4 parental and *PREX2*^KO^ lines. Right panel, frequency and location of indels upstream and downstream of the targeted cut site in *PREX2*^KO^ lines. B) Left panel, Sanger sequencing trace depicting region immediately upstream and downstream of the targeted PAM site in *PREX1* in WM266.4 parental and *PREX1*^KO^ lines. Right panel, frequency and location of indels upstream and downstream of the targeted cut site in *PREX1*^KO^ lines. C) Left panel, Sanger sequencing trace depicting region immediately upstream and downstream of the targeted PAM site in *PIK3CB* in WM266.4 parental and *PIK3CB*^KO^ lines. Right panel, frequency and location of indels upstream and downstream of the targeted cut site in *PIK3CB*^KO^ lines. In C–E, right panels, gene editing efficiency (%) as quantified using TIDE. D) Left panel, relative confluence of WM266.4 *PREX1*^KO^ cells treated with indicated treatments over time. Data, mean ± SEM (confluence relative to starting point). Right panel, relative change in confluence of WM266.4 *PREX1*^KO^ cells over indicated 96 h treatment. n=3 per treatment. Centre line, mean. p-values calculated by one-way ANOVA corrected for multiple comparisons (Tukey). E) Left panel, relative confluence of WM266.4 *PREX2*^KO^ cells treated with indicated treatments over time. Data, mean ± SEM (confluence relative to starting point). Right panel, relative change in confluence of WM266.4 *PREX2*^KO^ cells over indicated 96 h treatment. n=4 per treatment. Centre line, mean. p-values calculated by one-way ANOVA corrected for multiple comparisons (Tukey).

**Supplementary Fig. S8:**
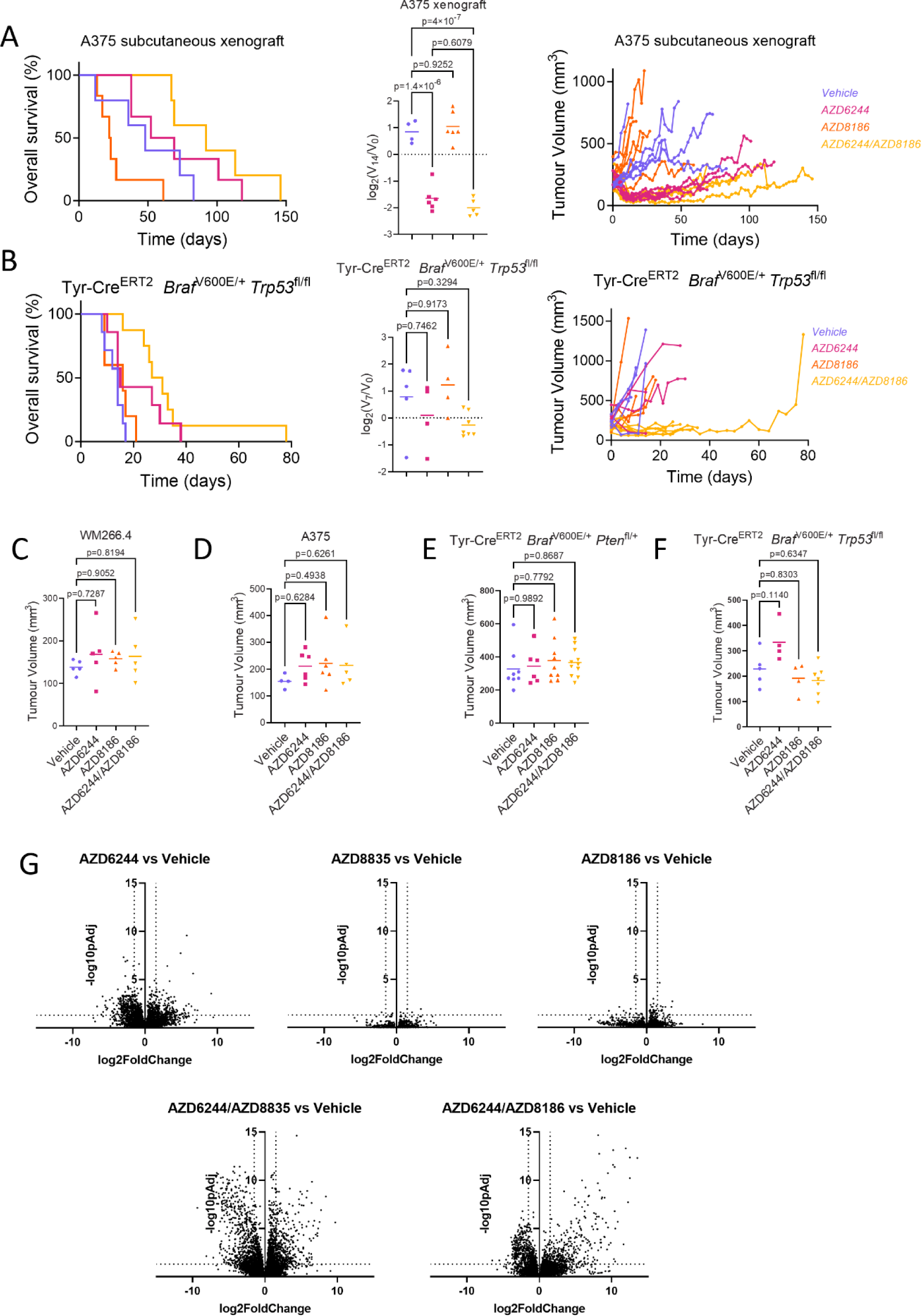
Co-targeting of MEK1/2 and p110β has therapeutic efficacy in melanoma *in vivo*. A) Left panel, Kaplan–Meier overall survival of mice harbouring A375 subcutaneous xenografts treated with vehicle (n=5; median survival, 48 days), AZD6244 (n=6; median survival, 60.5 days), AZD8186 (n=6; median survival, 22.5 days) or AZD6244/AZD8186 (n=5; median survival, 92 days). p-values calculated by log-rank (Mantel–Cox) test – vehicle vs AZD6244; p=0.3009, vehicle vs AZD8186; p=0.1130, vehicle vs AZD6244/AZD8186; p=0.0554 and AZD6244 vs AZD6244/AZD8186; p=0.3227. Centre panel, relative change in tumour volume of A375 xenografts over the first 7 days of indicated treatment. Vehicle (n=4), AZD6244 (n=6), AZD8186 (n=6), AZD6244/AZD8186 (n=5); Centre line represents mean. p-values calculated by one-way ANOVA corrected for multiple comparisons (Tukey). Right panel, longitudinal growth of individual tumours from mice harbouring A375 subcutaneous xenografts treated with vehicle (n=5) *vs* AZD6244 (n=6), AZD8186 (n=6), and AZD6244/AZD8186 (n=5). B) Left panel, Kaplan–Meier overall survival of BRAF P53 mice treated with vehicle (n=7; median survival, 14 days), AZD6244 (n=7; median survival, 15 days), AZD8186 (n=5; median survival, 16 days) or AZD6244/AZD8186 (n=8; median survival, 29 days). p-values were calculated by log-rank (Mantel–Cox) test – vehicle vs AZD6244, p=0.0869; vehicle vs AZD8186, p=0.2818; vehicle vs AZD6244/AZD8186, p=0.0002 and AZD6244 vs AZD6244/AZD8186, p=0.2318. Centre panel, relative change in BRAF P53 tumour volume over the first 7 days of indicated treatment. Vehicle (n=5), AZD6244 (n=4), AZD8186 (n=4), AZD6244/AZD8186 (n=7). Centre line represents mean. p-values calculated by one-way ANOVA corrected for multiple comparisons (Tukey). Right panel, longitudinal growth of individual tumours from vehicle-treated (n=5) *vs* AZD6244-treated (n=4), AZD8186-treated (n=4), and AZD6244/AZD8186-treated (n=7) BRAF P53 cohorts. C–F) Pre-treatment tumour volume in WM266.4 xenograft (C), A375 xenograft (D), BRAF PTEN (E), and BRAF P53 (F) cohorts. WM266.4: Vehicle, n=5; AZD6244, n=5; AZD8186, n=5; AZD6244/AZD8186, n=5. A375: Vehicle, n=4; AZD6244, n=6; AZD8186, n=6; AZD6244/AZD8186, n=5. BRAF PTEN: Vehicle, n=8; AZD6244, n=6; AZD8186, n=9; AZD6244/AZD8186, n=11. BRAF P53: Vehicle, n=5; AZD6244, n=4; AZD8186, n=4; AZD6244/AZD8186, n=7. Centre line represents mean. p-values calculated by one-way ANOVA corrected for multiple comparisons (Tukey). G) Volcano plots of log_2_FC *vs* -log_10_p_adj_ of transcripts from BRAF PTEN melanoma *in vivo* following 5-day treatment with AZD6244, AZD8835, AZD8186 and combinations thereof. Horizontal dashed lines represent linear p_adj_-value of 0.05, vertical dashed lines represent log2FC of 1.5.

